# Centromere drive and suppression by parallel pathways for recruiting microtubule destabilizers

**DOI:** 10.1101/2020.11.26.400515

**Authors:** Tomohiro Kumon, Jun Ma, Derek Stefanik, Erik C. Nordgren, R. Brian Akins, Junhyong Kim, Mia T. Levine, Michael A. Lampson

## Abstract

Selfish centromere DNA sequences bias their transmission to the egg in female meiosis. Evolutionary theory suggests that centromere proteins evolve to suppress costs of this “centromere drive”. In hybrid mouse models with genetically different maternal and paternal centromeres, selfish centromere DNA exploits a kinetochore pathway to recruit microtubule-destabilizing proteins that act as drive effectors. We show that such functional differences are suppressed by a parallel pathway for effector recruitment by heterochromatin, which is similar between centromeres in this system. Disrupting heterochromatin by CENP-B deletion amplifies functional differences between centromeres, whereas disrupting the kinetochore pathway with a divergent allele of CENP-C reduces the differences. Molecular evolution analyses using newly sequenced Murinae genomes identify adaptive evolution in proteins in both pathways. We propose that centromere proteins have recurrently evolved to minimize the kinetochore pathway, which is exploited by selfish DNA, relative to the heterochromatin pathway that equalizes centromeres, while maintaining essential functions.

## Introduction

Centromere evolution is paradoxical in that both repetitive centromere DNA and centromere-binding proteins evolve rapidly despite the conserved requirement of centromeres for faithful chromosome segregation (Henikoff et al., 2001; Lampson and Black, 2017; Malik and Henikoff, 2001; Melters et al., 2013; Schueler et al., 2010). Centromere DNA repeat monomer sequence and repeat copy number diverge between even closely related species. Repeat copy number also varies within species, for example, between human individuals or between mouse strains (Iwata-Otsubo et al., 2017; Langley et al., 2019). To explain this rapid evolution, the centromere drive hypothesis proposes that selfish centromere DNA sequences (either monomer sequence variants or repeat number expansions) drive in female meiosis by increasing their transmission rate to the egg. Potential deleterious consequences of driving centromeres, such as meiotic segregation errors, would select for centromere-binding protein variants that suppress these fitness costs (Finseth et al., 2020; Fishman and Saunders, 2008; Henikoff et al., 2001). New selfish DNA variants subsequently arise to start another cycle of drive and suppression in a continual evolutionary arms race.

Our previous work leveraged natural variation in mouse centromere DNA to study the molecular mechanisms of centromere drive (Akera et al., 2017, 2019; Chmátal et al., 2014; Iwata-Otsubo et al., 2017). Selfish centromeres in these model systems recruit more effector proteins that destabilize interactions with spindle microtubules, allowing them to detach from microtubules that would otherwise direct them to the polar body. Microtubule detachment and reattachment reorients the selfish centromeres toward the egg side of the meiosis I spindle (Akera et al., 2019) (Figure 1A). This reorientation depends on BUB1 kinase at kinetochores, which phosphorylates pericentromeric histone H2A. Phosphorylated H2A recruits Shugoshin-2 (SGO2), which recruits microtubule destabilizing proteins such as MCAK (Akera et al., 2019) (Figure 1B, kinetochore pathway). In one intra-species *Mus musculus domesticus* hybrid, selfish centromeres with expanded minor satellite DNA repeats assemble more centromere chromatin containing the histone H3 variant CENP-A (Iwata-Otsubo et al., 2017). These expanded centromeres also form larger kinetochores with more BUB1 kinase, leading to more effectors (SGO2 and MCAK) (Akera et al., 2019). In this hybrid, the larger centromeres are from a standard laboratory strain (either CF-1 or C57BL/6J), which is crossed to a wild-derived strain (CHPO) with smaller centromeres. Thus, the centromeres of paired homologous chromosomes within a meiotic bivalent are both genetically and functionally different in the hybrid (Figure 1C). These findings show how selfish centromeres can drive by recruiting more effectors. How centromere-binding proteins can evolve to suppress the costs of drive remains an open question despite being a crucial component of the centromere drive model. Details of the fitness costs are unclear, but they likely depend on functional differences between paired centromeres in meiosis and would therefore be suppressed by reducing these differences.

**Figure 1:**
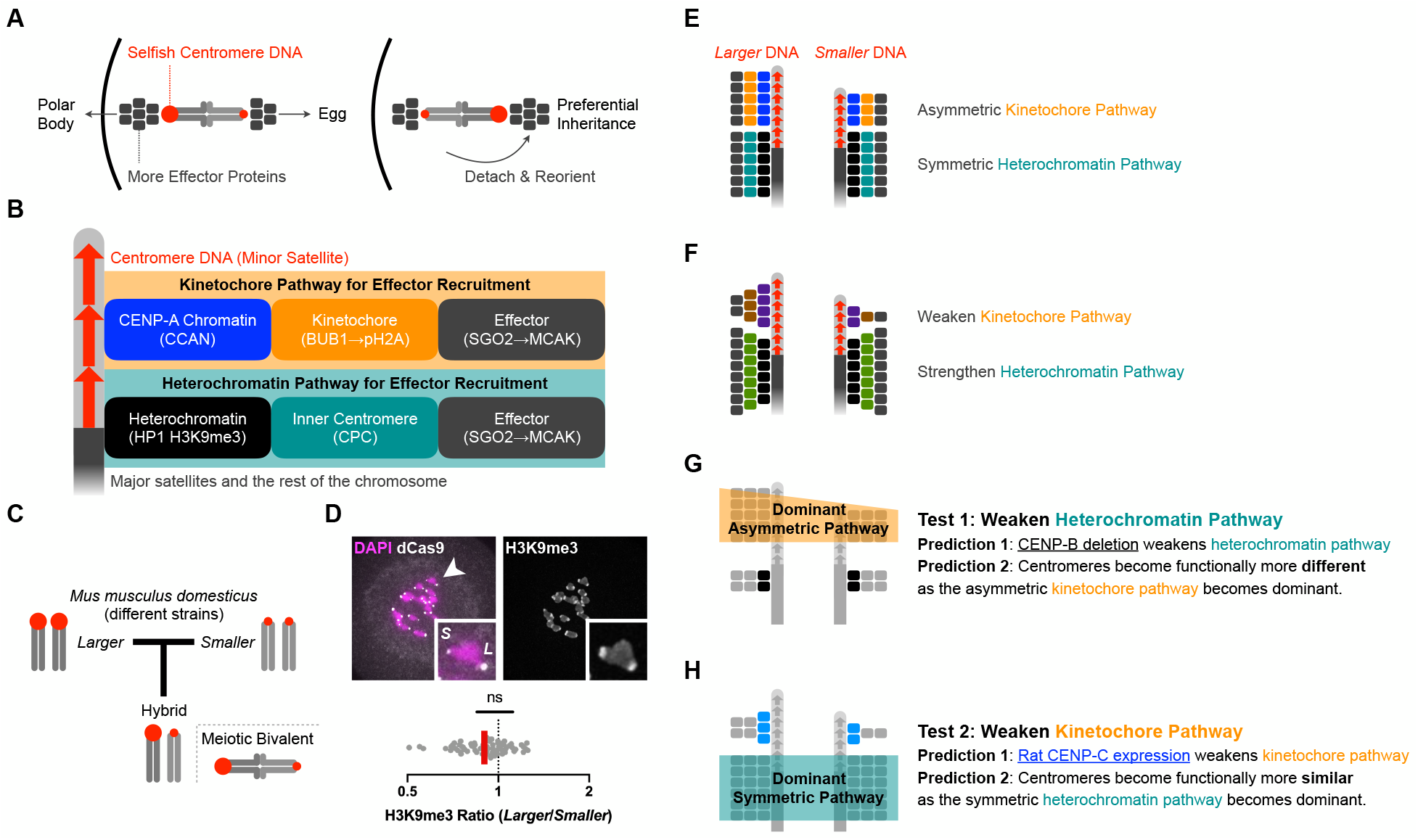
Parallel pathway model for centromere drive and suppression. (A) Centromere drive by recruiting effector proteins that destabilize interactions with spindle microtubules. Selfish centromeres recruit more effector proteins (SGO2 and MCAK), preferentially detach from microtubules when facing the cortical side of the spindle, and reorient to bias their segregation to the egg. The cortical side of a meitoic bivalent will segregate to the polar body, whereas the other side will segregate to the egg. (B) Two pathways for effector recruitment. CENP-A and the CCAN (constitutive centromere-associated network) connect centromere DNA to the kinetochore, which assembles during meiosis or mitosis. Kinetochore-localized BUB1 kinase phosphorylates pericentromeric histone H2A to recruit SGO2. In parallel, pericentromeric heterochromatin also recruits SGO2 via the CPC (chromosome passenger complex) at the inner centromere. Although the two pathways are shown as independent here for simplicity, they are not completely separate, as the CPC is also recruited by SGO2 via the kinetochore pathway (Yamagishi et al., 2010). This contribution to effector recruitment depends on the kinetochore, so we consider it part of the kinetochore pathway. (C) CHPO hybrid model system. Crossing strains with larger (CF-1) and smaller (CHPO) centromeres generates a hybrid in which genetically different centromeres are paired in meiotic bivalents. Larger red circles indicate more minor satellite centromere DNA repeats. (D) CHPO hybrid oocytes were microinjected with cRNA for dCas9-GFP and gRNA targeting minor satellite centromere DNA to distinguish larger (*L*) and smaller (*S*) centromeres, fixed at meiosis I, and stained for H3K9me3. The H3K9me3 ratio for each pair of larger and smaller centromeres within a bivalent is plotted (n=67 bivalents); red line, geometric mean; ns: no significant deviation from 1. (E) Symmetric heterochromatin pathway and asymmetric kinetochore pathway in our hybrid model system. The amount of recruited proteins is represented by the number of boxes, using the same color code as panel B. (F) Parallel pathway model for suppression of functional differences between centromeres by recruiting similar amounts of effector proteins on genetically different centromeres. Colored boxes represent changes relative to panel E. Proteins in the kinetochore pathway can adapt by reducing affinity for DNA or for other proteins leading to effector recruitment. Inner centromere proteins can adapt by increasing affinity for heterochromatin or for effectors. (G) As CENP-B recruits heterochromatin proteins, deleting CENP-B weakens the heterochromatin pathway for effector recruitment (prediction 1), making the asymmetric kinetochore pathway dominant and centromeres functionally more asymmetric (prediction 2). (H) Introducing a divergent allele of CENP-C (blue boxes) disrupts interactions for effector recruitment and therefore weakens the kinetochore pathway (prediction 1) and makes centromeres functionally more similar (prediction 2).

### The parallel pathway model for drive and suppression provides three testable predictions

Based on our finding that selfish centromeres drive by recruiting more effectors, we propose that functional differences between centromeres can be suppressed by equalizing effector recruitment via a second pathway. This equalization would render genetically different centromeres functionally equivalent. This model incorporates previous findings that in addition to the kinetochore pathway, which acts through BUB1 kinase, effectors are also recruited through a heterochromatin pathway. Pericentromeric heterochromatin recruits the chromosome passenger complex (CPC), which recruits SGO2 and MCAK (Figure 1B, heterochromatin pathway) (Abe et al., 2016; Ainsztein et al., 1998; Tsukahara et al., 2010). In our CHPO hybrid model system (Figure 1C), the kinetochore pathway is asymmetric: we observe higher levels of the kinetochore proteins HEC1/NDC80 and CENP-C on larger vs smaller centromeres (Chmátal et al., 2014; Iwata-Otsubo et al., 2017). In contrast, the heterochromatin pathway is symmetric: the heterochromatin mark, H3K9me3, is equal on the two sides of each bivalent (Figure 1D and 1E) (Iwata-Otsubo et al., 2017). These observations suggest that, in this system, selfish centromere DNA exploits the kinetochore pathway to make genetically different centromeres also functionally different, with larger centromeres recruiting more effectors. In contrast, the heterochromatin pathway appears insensitive to selfish DNA, recruiting effectors equally. We propose that the host evolves to suppress functional differences by centromere protein innovations that minimize the contribution of the asymmetric kinetochore pathway to effector recruitment, relative to the symmetric heterochromatin pathway.

Evolution of the kinetochore pathway is constrained by its indispensable role in mitotic and meiotic chromosome segregation. Nevertheless, proteins may evolve to weaken the pathway by reducing interactions between centromere-binding proteins and DNA or between proteins leading to effector recruitment (Figure 1F). Similarly, evolution of heterochromatin proteins is constrained by numerous vital heterochromatin-dependent cellular functions (Allshire and Madhani, 2017). Inner centromere proteins (such as the CPC) that interact with heterochromatin may evolve, however, to increase effector recruitment. Finally, overall effector levels are also constrained because microtubule destabilizing activity is necessary to correct kinetochore-microtubule attachment errors, but excessive destabilizing activity weakens attachments necessary for anaphase segregation and activates the spindle assembly checkpoint (Godek et al., 2014). According to our parallel pathway model, a new centromere DNA variant can exploit the kinetochore pathway to recruit more effectors by strengthening interactions with any centromere-binding protein that contacts the DNA: CENP-A, the CENP-A chromatin assembly machinery, or other proteins that link centromere chromatin to the kinetochore (e.g., CENP-C or CENP-T). To suppress functional differences between centromeres, the centromere protein network recurrently evolves to minimize the kinetochore pathway relative to the heterochromatin pathway while maintaining essential functions.

Here we test three predictions from the parallel pathway model. First, when the symmetric heterochromatin pathway is weakened, we predict that the asymmetric kinetochore pathway makes a relatively larger contribution to effector recruitment. Genetically different centromeres in our hybrid model system should therefore become functionally more different. To target pericentromeric heterochromatin, we deleted CENP-B, which is the only centromeric chromatin component that is dispensable for core centromere function. CENP-B is recently acquired in mammals and fission yeast from a pogo-like transposase (Casola et al., 2007; Kipling and Warburton, 1997), and several domesticated transposases regulate heterochromatin (Gao et al., 2020; Jangam et al., 2017; Nozawa et al., 2010; Yang et al., 2017). In mouse and human cultured cells and fission yeast, CENP-B contributes to pericentromeric heterochromatin formation via heterochromatin protein recruitment (Nakagawa et al., 2002; Okada et al., 2007; Otake et al., 2020), so deleting CENP-B should weaken the heterochromatin pathway (Figure 1G). Mammalian CENP-B can also contribute to the kinetochore pathway via CENP-C recruitment (Fachinetti et al., 2015), so the functional consequences of CENP-B deletion in our model need to be tested. Second, when the asymmetric kinetochore pathway is weakened, we predict that centromeres become functionally more similar due to the symmetric heterochromatin pathway. We selected CENP-C as a key scaffold protein in the kinetochore pathway that is known to evolve rapidly under positive selection (Klare et al., 2015; Schueler et al., 2010; Talbert et al., 2004). Under the parallel pathway model, CENP-C interfaces have co-evolved with interacting partners to modulate effector recruitment. Thus, introducing a divergent allele of CENP-C in mouse cells (e.g., rat CENP-C, in which 32% of the amino acid sequence is different) is predicted to disrupt such interactions and weaken the kinetochore pathway (Figure 1H). Third, if proteins in the kinetochore and heterochromatin pathways have evolved to modulate effector recruitment, we predict signatures of positive selection in multiple protein domains involved in effector recruitment. In contrast, the previous model of an arms race limited to interactions between centromere DNA and DNA-binding proteins only predicts rapid evolution of protein domains involved in DNA binding (Henikoff et al., 2001; Malik and Henikoff, 2001). Our observations are consistent with all three predictions, supporting our parallel pathway model for drive and suppression.

### Deleting CENP-B weakens the heterochromatin pathway and makes centromeres functionally more asymmetric

To determine the contribution of CENP-B to effector recruitment, we created *Cenpb* null mice using CRISPR genome editing (Supplementary Figure 1A-1C). We find that loss of CENP-B weakens both the kinetochore and heterochromatin pathways, as shown by reduced CENP-C and H3K9me3 staining, respectively (Figure 2A). These results are consistent with previous findings that CENP-B contributes to CENP-C recruitment and to formation of pericentromeric heterochromatin (Fachinetti et al., 2015; Okada et al., 2007; Otake et al., 2020). We also find reduced effector recruitment, as represented by SGO2 staining (Figure 2A), consistent with the idea that CENP-B recruits effectors through the kinetochore and heterochromatin pathways.

**Figure 2:**
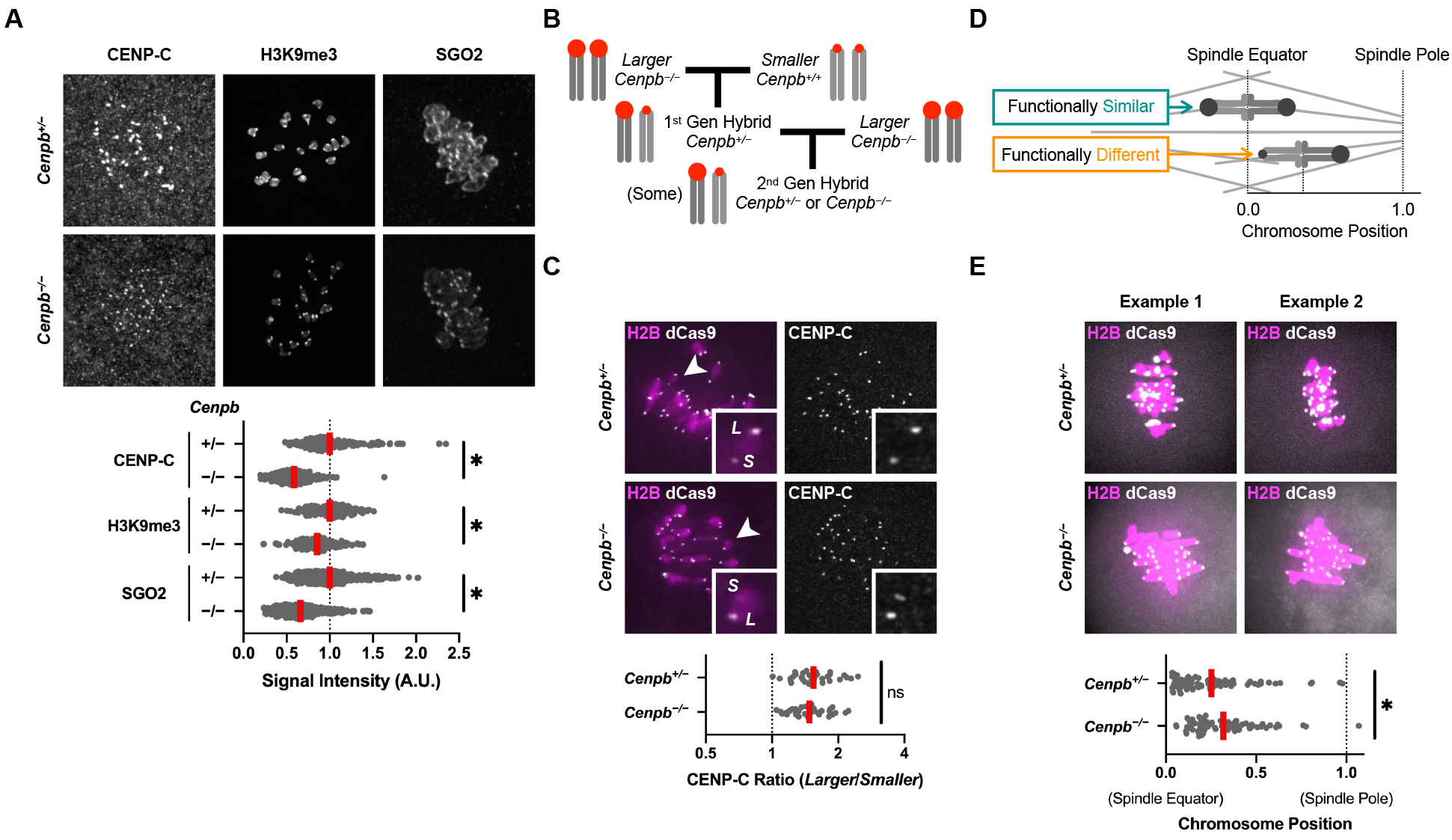
Deleting CENP-B weakens the heterochromatin pathway and makes centromeres functionally more asymmetric. (A) *Cenpb*^*+/−*^ or *Cenpb*^*−/−*^ oocytes were fixed in meiosis I and stained for CENP-C, H3K9me3, or SGO2. Plot shows centromere signal intensities, normalized by mean intensity of *Cenpb*^*+/−*^ control for each protein. Each dot represents a single centromere (n≥154 centromeres for each condition); red line, mean; *p<0.05. (B) Crossing scheme to produce second generation hybrid *Cenpb*^*−/−*^ mice with larger and smaller paired centromeres. (C) Second generation hybrid oocytes were microinjected with cRNA for dCas9-EGFP and gRNA targeting minor satellite centromere DNA, fixed in meiosis I, and stained for CENP-C. The CENP-C ratio for each pair of larger (*L*) and smaller (*S*) centromeres on a bivalent is plotted (n=34 bivalents for each genotype); red line, geometric mean; ns: not significant. (D) Schematic of chromosome position assay to measure functional differences between paired centromeres. Distance from the spindle pole to the equator is defined as 1 for each cell to normalize for variation in spindle size. (E) Second generation hybrid oocytes were microinjected with cRNAs for dCas9 and H2B and gRNA targeting minor satellite centromere DNA. Cells were imaged live to preserve chromosome positions, measured at late metaphase I. In the plot, each dot represents a single bivalent (n≥74 bivalents for each genotype); red line, mean.

The known functions of CENP-B suggest two hypotheses for how it might affect centromeres in our CHPO hybrid model system. First, as the only centromere protein known to recognize a specific DNA sequence (the CENP-B box in repetitive centromere DNA) (Masumoto et al., 1989), CENP-B could be exploited by selfish larger centromeres with more CENP-B boxes to increase asymmetry via the kinetochore pathway. Second, CENP-B may suppress functional differences between centromeres by increasing the symmetric heterochromatin pathway. To test these hypotheses, we generated *Cenpb* null mice with paired larger and smaller centromeres through two generations of crosses (Figure 2B and Supplementary Figure 1D), and analyzed kinetochore pathway asymmetry and functional differences between centromeres. To determine the impact of CENP-B on the kinetochore pathway, we analyzed CENP-C in meiotic bivalents with paired larger and smaller centromeres in second-generation hybrid *Cenpb*^*−/−*^ oocytes. CENP-C was reduced to a similar extent on both larger and smaller centromeres (Supplementary Figure 1E) and consistent with this equivalent reduction, the kinetochore asymmetry remained intact (Figure 2C). Therefore, CENP-B does not contribute to asymmetry in the kinetochore pathway, arguing against the first hypothesis that selfish centromere DNA exploits the kinetochore pathway via CENP-B.

To test the second hypothesis, that CENP-B acts as a suppressor through the symmetric heterochromatin pathway (Figure 1G), we examined functional differences between centromeres in second-generation hybrid oocytes. As a readout, we analyzed chromosome position on the spindle at metaphase I (Figure 2D). Chromosome position is sensitive to differences in interactions with spindle microtubules between centromeres of homologous chromosomes, which are paired in a meiotic bivalent. If the paired centromeres are genetically and functionally similar, then chromosomes align at the spindle equator in a typical metaphase configuration. In our hybrid model systems, paired centromeres are genetically and functionally different, and bivalents are positioned off-center on the spindle, with the larger centromere closer to its attached pole (Akera et al., 2019; Chmátal et al., 2014). Manipulations that make these genetically different centromeres functionally more similar will lead to positioning closer to the spindle equator, as previously shown by manipulating BUB1 kinase to equalize MCAK levels on larger and smaller centromeres (Akera et al., 2019). Conversely, manipulations that make the centromeres functionally more different will position bivalents closer to the poles. We find that asymmetric bivalents with genetically different centromeres are positioned more off-center, closer to the spindle poles, in *Cenpb*^*−/−*^ compared to control *Cenpb*^*+/−*^ oocytes (Figure 2E). This result indicates that paired larger and smaller centromeres are functionally more different in the absence of CENP-B, consistent with the prediction that the symmetric heterochromatin pathway is weakened, making the asymmetric kinetochore pathway relatively more dominant (Figure 1G).

### Introducing a divergent CENP-C allele weakens the kinetochore pathway and makes centromeres functionally more symmetric

To weaken the kinetochore pathway, we targeted CENP-C because it serves as a hub for recruiting kinetochore proteins. Because it is an essential gene, we introduced a divergent allele rather than deleting it. Our model predicts that CENP-C has co-evolved with interacting partners to modulate effector recruitment, so that an allele from another species will disrupt these interactions and weaken the kinetochore pathway (Figure 1H, Prediction 1). To test this prediction, we selected rat as a model organism close to mouse with divergent centromere DNA and proteins (Gibbs et al., 2004; Takeiri et al., 2013). Because protein interfaces change by genetic drift as well as by selection, an allele from a closely related species minimizes incompatibilities coming from stochastic changes. We find that effector recruitment, as represented by SGO2 staining, is reduced when rat CENP-C is expressed in mouse oocytes, compared to mouse CENP-C (Figure 3A). This result is consistent with our model prediction and could reflect differences between mouse and rat CENP-C in their recruitment to centromeres or in their interactions with other kinetochore proteins. For example, evolution at an interface with CENP-A nucleosomes or with CENP-B may disrupt rat CENP-C recruitment to centromeres. Alternatively, CENP-C evolution might impact the domains that mediate interactions with other kinetochore proteins involved in SGO2 recruitment. We find that mouse and rat CENP-C are equally recruited to mouse centromeres (Figure 3B), indicating functional changes at an interface with other kinetochore proteins.

**Figure 3:**
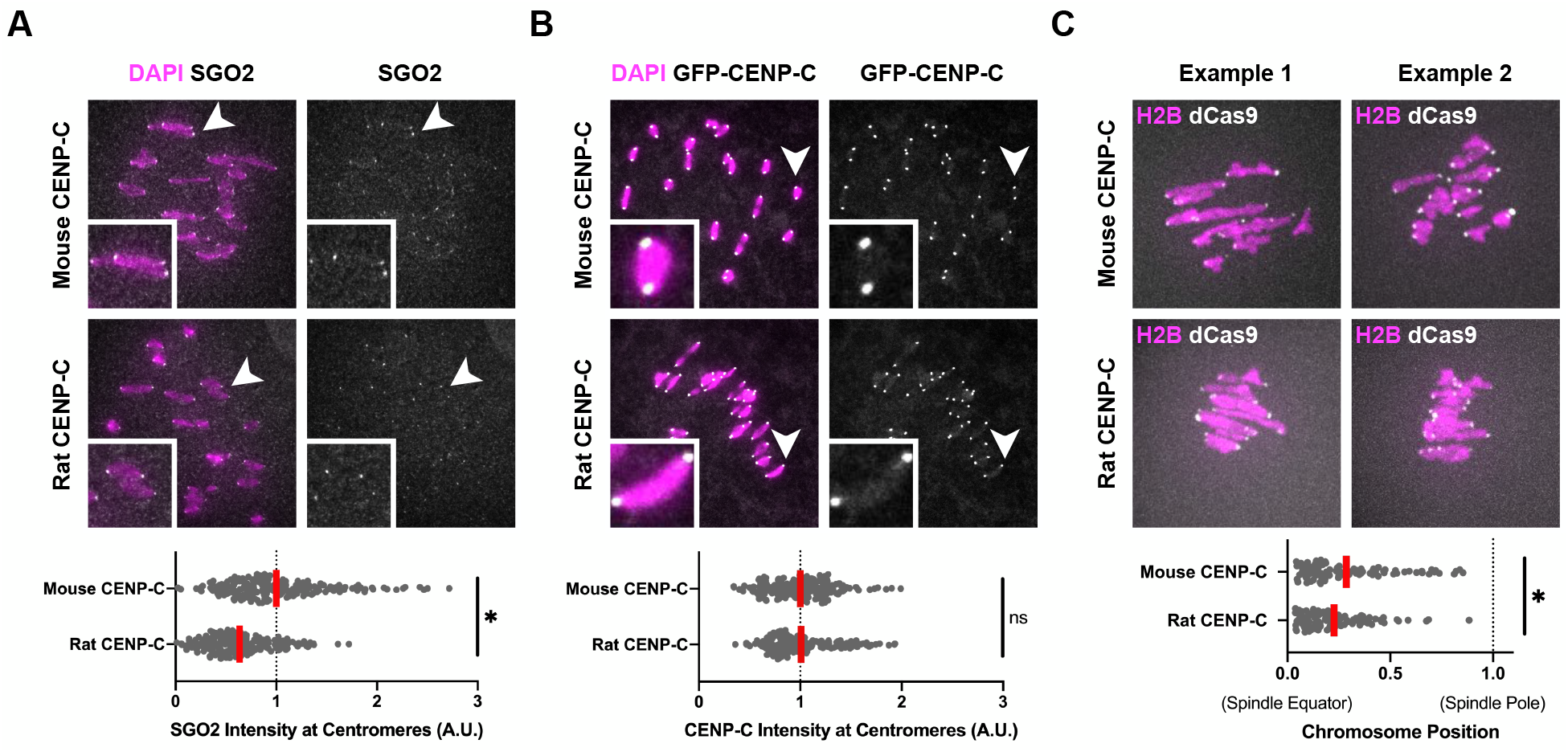
Introducing a divergent CENP-C allele weakens the kinetochore pathway and makes centromeres functionally more symmetric. (A and B) CF-1 oocytes were microinjected with cRNA for GFP-tagged mouse or rat CENP-C and fixed in meiosis I. Cells were stained for SGO2 (A) or analyzed for GFP fluorescence (B). Plots show centromere signal intensities. Each dot represents a single centromere (n=200 centromeres from 20 oocytes for each construct); red line, mean; *p<0.05; ns: not significant. (C) CHPO hybrid oocytes (see Figure 1C) were microinjected with cRNA for GFP-tagged mouse or rat CENP-C, together with cRNAs for GFP-tagged H2B and mCherry-tagged dCas9 and gRNA targeting minor satellite centromere DNA. Cells were imaged live to preserve chromosome positions, measured at late metaphase I. In the plot, each dot represents a single bivalent (n=100 bivalents from ≥20 oocytes for each construct); red line, mean.

Rat CENP-C expression provides an experimental tool to weaken the kinetochore pathway, allowing us to test our prediction that genetically different centromeres become functionally more similar in our hybrid model system (Figure 1H, Prediction 2). Using chromosome position as a functional readout of centromere asymmetry (Figure 2D), we find that expression of rat CENP-C in CHPO hybrid oocytes (Figure 1C) leads to bivalents positioned closer to the spindle equator (Figure 3C). This result indicates that the paired larger and smaller centromeres are functionally more similar, consistent with the prediction that the symmetric heterochromatin pathway becomes relatively more dominant when the asymmetric kinetochore pathway is weakened (Figure 1H, Prediction 2).

### Proteins in the kinetochore and heterochromatin pathways have signatures of recurrent adaptive evolution

The original model of centromere drive and suppression posits an arms race between selfish centromere DNA and DNA-binding proteins such as CENP-A (Henikoff et al., 2001; Malik and Henikoff, 2001). This model predicts adaptive evolution of centromere protein domains that physically interact with DNA, and conservation of domains and other centromere proteins that do not bind DNA. In contrast, our parallel pathway model predicts signatures of recurrent adaptive evolution in protein domains leading to effector recruitment, including those that do not directly contact centromere DNA (Figure 4A). These changes could either weaken the kinetochore pathway or strengthen the heterochromatin pathway to make genetically different centromeres functionally more similar (Figure 1F). Rapid evolution of centromere proteins has been reported in several eukaryotic lineages, but there are no mechanistic studies of drive in these lineages (Finseth et al., 2015; Malik and Henikoff, 2001; Schueler et al., 2010; van der Lee et al., 2017). To analyze centromere protein evolution in a system where we have identified drive effectors, we tested for signatures of positive selection in Murinae. Because the sparseness of the phylogenetic tree of currently available Murinae genomes limits our statistical power to detect positive selection, we sequenced six new genomes (Figure 4B) using linked-read whole genome sequencing (10x Genomics). Each genome was assembled onto the *Mus musculus* reference genome (mm10) with LongRanger and *de novo* assembled with Supernova (see Methods and Supplementary Table 1). Sampling evolutionary time more comprehensively increases our opportunities to observe adaptive changes (and minimize false positives from stochastic changes by genetic drift), especially those adaptive changes that are common to multiple independent lineages. Thus, these genomes provide a valuable resource for molecular evolution approaches in mouse as a mammalian model organism, such as our analyses of centromere proteins discussed below.

**Figure 4:**
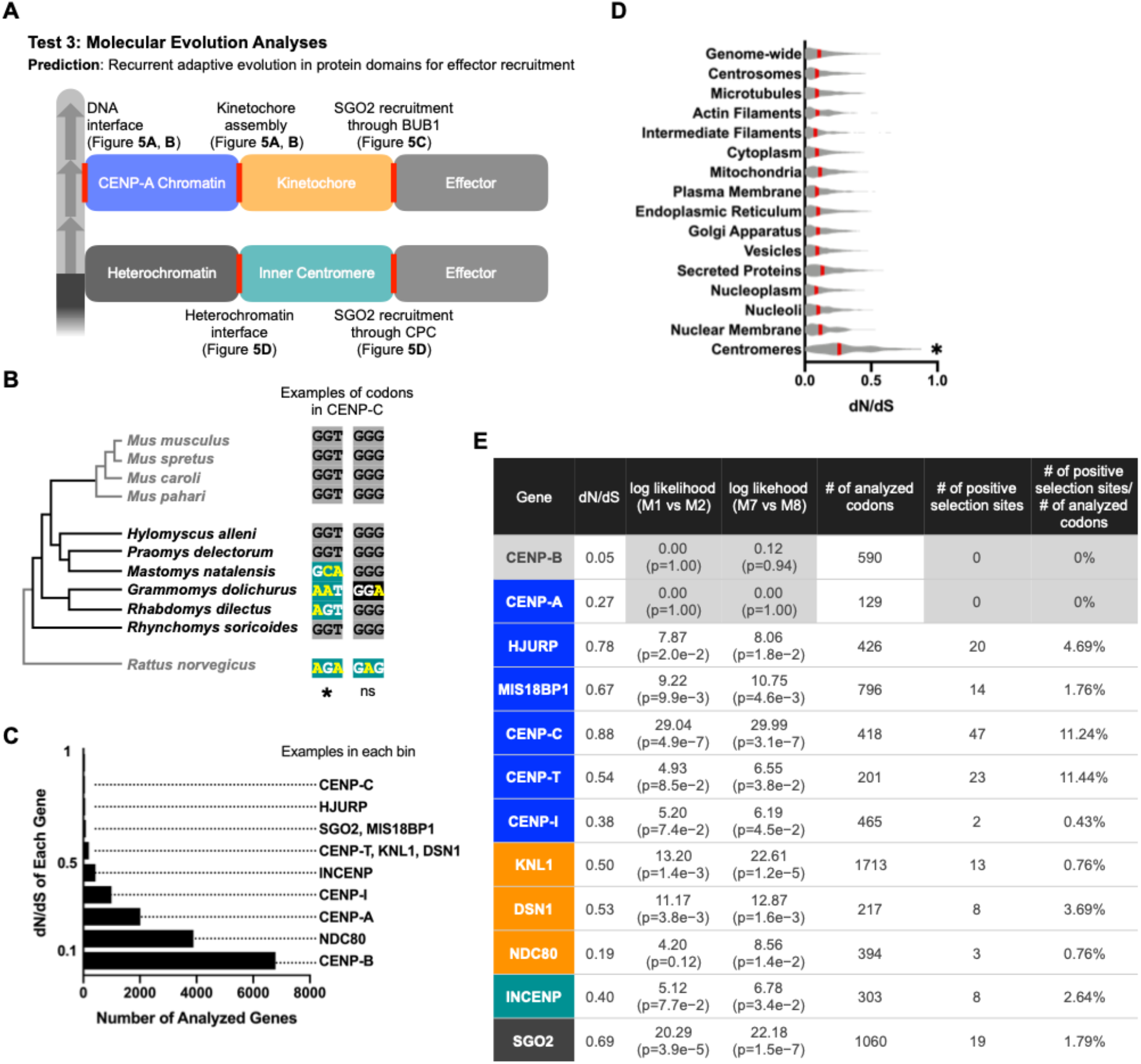
Proteins in the kinetochore and heterochromatin pathways have signatures of recurrent adaptive evolution. (A) Our parallel pathway model predicts that proteins in both pathways will have signatures of recurrent adaptive evolution at interfaces (shown in red) that lead to effector recruitment. (B) Phylogenetic tree of Murinae species shows previously available genomes in gray and our newly sequenced genomes in black. Example codons show positive selection or neutral changes (mouse CENP-C Gly469 and Gly470). Nucleotide substitutions are shown in yellow, with synonymous and nonsynonymous substitutions highlighted in black or green, respectively. Higher numbers of nonsynonymous substitutions are interpreted as adaptive change under positive selection. PAML analysis: *P>0.99 for positive selection or not significant (ns) indicating a neutral change. (C) Histogram shows the number of genes in each bin of dN/dS values, with examples of genes in each bin. (D) Average dN/dS across subcellular compartments. Red line, mean; *p<0.05 for comparison to all other compartments. (E) To test for signatures of positive selection in PAML, the likelihood of models of neutral codon evolution (M1 or M7) are compared to models allowing positive selection (M2 or M8). CENP-A and CENP-B are examples of genes without signatures of positive selection (see Supplementary Table 2 for other genes and Supplementary Figure 2 for further analyses of CENP-A and CENP-B). The number of analyzed codons is less than the total protein length as insertions, deletions, and ambiguous alignment are not analyzed. The number of positive selection sites is the number of codons with P>0.90 from Naive Empirical Bayes (NEB) analysis or Bayes Empirical Bayes (BEB) analysis from model 2 or 8.

Low rates of nonsynonymous substitutions, which change the encoded amino acid, relative to synonymous substitutions (dN/dS) indicate purifying selection, as deleterious substitutions are selected against. Higher dN/dS indicates either adaptive evolution or loss of constraint, necessitating further analysis to identify signatures of positive selection (Echave et al., 2016; Sironi et al., 2015). We calculated dN/dS for all annotated mouse-rat orthologous genes. We find that multiple genes encoding centromere proteins have high dN/dS relative to the genome overall (Figure 4C), and the average dN/dS for these genes is significantly higher than for any other subcellular compartment (Figure 4D). We selected 46 genes with well-characterized centromere functions to analyze for signatures of positive selection based on phylogenetic analysis, using PAML (Yang, 2007). Consistent with our prediction, we find such signatures at genes in multiple modules of the kinetochore and heterochromatin pathways (Figure 4E).

Extensive previous studies of centromere organization and function have established functional modules which can recruit drive effectors either directly or indirectly (Figure 4A). To fit our observations into this framework, we assigned genes to these modules. One module is CENP-A chromatin and the constitutive centromere-associated network (CCAN). Selfish centromere DNA can increase effector recruitment by expanding CENP-A chromatin through increased deposition of CENP-A nucleosomes. This process depends on a specialized histone chaperone, HJURP, which is targeted to centromeres by the MIS18 complex though interactions with CENP-C or CENP-I (Dunleavy et al., 2009; Foltz et al., 2009; Fujita et al., 2007; Moree et al., 2011; Shono et al., 2015). We find rapid evolution of HJURP, MIS18BP1, CENP-I, and the domain of CENP-C that interacts with the MIS18 complex (Figure 4E and 5A). In contrast, heterochromatin proteins such as HP1 paralogs and SUV39H1, which are not specific to centromeres/pericentromeres, are highly conserved (Supplementary Table 2), consistent with the idea that heterochromatin broadly suppresses selfish genetic elements regardless of the underlying DNA sequence (Allshire and Madhani, 2017). These findings suggest that selection acts on the CENP-A chromatin assembly pathway to prevent expansion, but selfish centromere DNA does not exploit the heterochromatin pathway, consistent with our observation that genetically different centromeres have symmetric heterochromatin in our intra-species and inter-species hybrids (Figure 1D and our unpublished data).

Under our model (Figure 1E), selfish centromere DNA can also recruit more effectors through the kinetochore pathway by strengthening direct interactions with CENP-A or with CCAN components, leading to larger kinetochores and more BUB1 kinase. Proteins can subsequently adapt by weakening interactions either with DNA or with other kinetochore proteins (Figure 4A, DNA interface and kinetochore assembly). Within the CCAN, CENP-C and CENP-T sub-pathways connect CENP-A chromatin to kinetochore proteins. The middle part of CENP-C interacts with CENP-A nucleosomes, while the N-terminus interacts with the MIS12 kinetochore complex (Petrovic et al., 2016; Weir et al., 2016). Similarly, the CENP-TWSX nucleosome-like complex contacts centromere DNA, and the other end of CENP-T interact with MIS12 and NDC80 kinetochore complexes (Cortes-Silva et al., 2020; Nishino et al., 2012; Veld et al., 2016). Consistent with our model, we detect signatures of positive selection in the chromatin-interacting domains and the kinetochore-interacting domains of both CENP-C and CENP-T (Figure 5A and 5B). In contrast, the DNA-interacting domain of CENP-B is conserved, consistent with our finding that selfish centromere DNA does not exploit CENP-B. Unlike in other eukaryotic lineages such as monkeyflower, fly, and primates (Finseth et al., 2015; Malik and Henikoff, 2001; Schueler et al., 2010), we do not detect signatures of positive selection in the part of CENP-A that can be aligned in Murinae species, but the N-terminal tail is duplicated in some species and therefore difficult to analyze by standard methods (Supplementary Figure 2A). Diversification of the CENP-A N-terminal tail is also observed in plants, where crosses between strains expressing different alleles exhibit zygotic segregation errors and genome elimination (Maheshwari et al., 2015).

**Figure 5:**
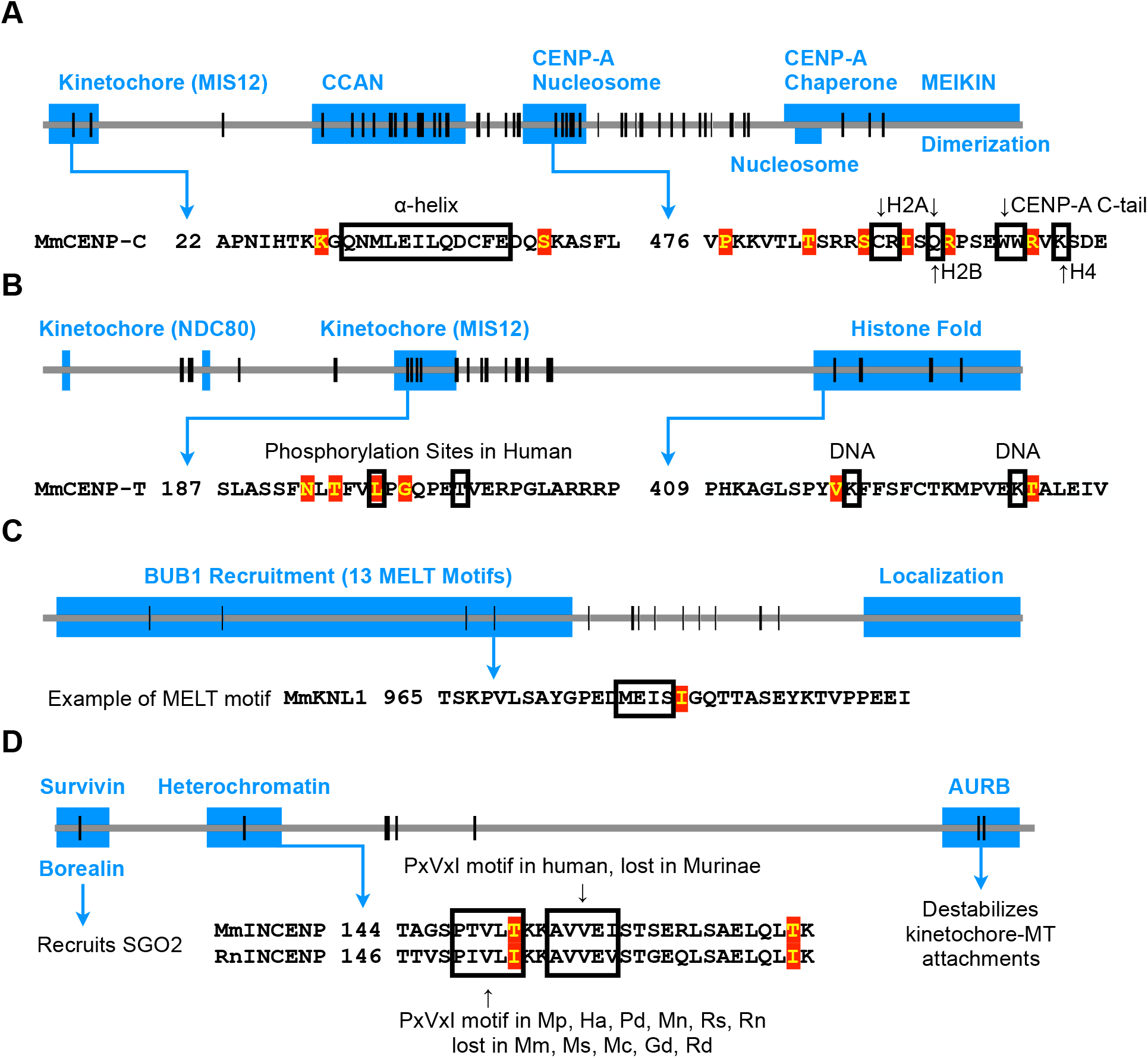
Protein domains that lead to microtubule destabilizer recruitment are recurrently evolved. Each horizontal line represents the entire protein for each gene, and vertical lines represent positions of positively selected amino acid residues. Blue boxes show known functional domains from previous studies. Amino acid sequences within domains of interest are shown, with positively selected residues highlighted in red and known functional residues outlined in black. (A) Signatures of positive selection are found throughout CENP-C. In the kinetochore domain, the a-helix interacts with MIS12 (Petrovic et al., 2016). The CCAN domain (also known as PEST domain) interacts with CENP-HIKM (Klare et al., 2015) and CENP-LN (Pentakota et al., 2017), and together forms the CENP-ACHIKMLN complex (Weir et al., 2016). In the domain interacting with CENP-A nucleosomes (also known as central region), residues interacting with H2A, H2B, H4 and the CENP-A C-terminal tail are indicated. This domain binds CENP-A nucleosomes more specifically than the more C-terminal nucleosome binding domain (also known as CENP-C motif), which also interacts with H3 nucleosomes (Allu et al., 2019; Kato et al., 2013). The CENP-C C-terminus has multiple functions, including M18BP1 recruitment (Dambacher et al., 2012), MEIKIN recruitment (Kim et al., 2015), and dimerization (Sugimoto et al., 1997). (B) Signatures of positive selection are found in the kinetochore interaction domain and histone fold domain of CENP-T. CDK1-dependent phosphorylation at Thr195 and Ser201 in human CENP-T (substituted with Leu and Thr, respectively, in mice) regulates MIS12 recruitment (Rago et al., 2015; Veld et al., 2016). Signatures of positive selection are detected around these regulatory residues for MIS12 recruitment. Some DNA interacting residues within the histone fold domain are shown (Nishino et al., 2012). (C) Signatures of positive selection are found in the domain of KNL1 that recruits BUB1 via repeated MELT motifs (Krenn et al., 2013). One MELT motif is shown as an example. (D) Signatures of positive selection are found in domains of INCENP that interact with Borealin/Survivin, with heterochromatin, and with Aurora B kinase. Heterochromatin recruits INCENP, and Borealin mediates the interaction with SGO2 (Abe et al., 2016; Ainsztein et al., 1998; Tsukahara et al., 2010). Survivin binds cohesin and pH3T3 at pericentromeres (Kelly et al., 2010; Wang et al., 2010; Yamagishi et al., 2010), providing another mechanism to localize the CPC. A PxVxI motif, which interacts with the HP1 chromoshadow domain, is present in some Murinae species and lost in others, shown with *Mus musculus* (Mm) and *Rattus norvegicus* (Rn) as examples. Other species from the phylogenetic tree in Figure 4B: *Mus spretus* (Ms), *Mus caroli* (Mc), *Mus pahari* (Mp), *Hylomyscus alleni* (Ha), *Praomys delectorum* (Pd), *Mastomys natalensis* (Mn), *Grammomys dolichurus* (Gd), *Rhabdomys dilectus* (Rd), and *Rhynchomys soricoides* (Rs).

In the kinetochore module, proteins can adapt to weaken the kinetochore pathway by reducing either kinetochore assembly or BUB1 binding to the kinetochore (Figure 4A). We find rapid evolution of the kinetochore proteins DSN1, KNL1, and NDC80. DSN1 is a component of the MIS12 complex which assembles onto the CCAN and serves as a platform for binding KNL1 and the NDC80 complex (Petrovic et al., 2014). KNL1 contains multiple protein docking motifs, including repeated MELT motifs that recruit BUB1 kinase (Musacchio and Desai, 2017). Thus, changes in DSN1 and KNL1 can regulate kinetochore assembly and BUB1 recruitment.

Consistent with the possibility that these interfaces evolve to modulate effector recruitment, we find signatures of positive selection in the MELT motifs of KNL1 (Figure 5C). NDC80 is the major microtubule binding protein in the kinetochore, but we find signatures of positive selection in the coiled-coil domain and not in the microtubule interacting domain. The coiled-coil domain recruits the SKA complex, which stabilizes kinetochore-microtubule attachment (Veld et al., 2019) and could be involved in counteracting destabilizing activities exploited by selfish centromeres.

Although selfish centromere DNA is likely unable to exploit heterochromatin to drive, components of the heterochromatin pathway, particularly inner centromere proteins, can adapt to increase effector recruitment relative to the kinetochore pathway in our model. In the inner centromere module, which links heterochromatin to effectors (Figure 4A), INCENP is a scaffold component of the CPC that interacts directly with heterochromatin and indirectly with SGO2 (Abe et al., 2016; Ainsztein et al., 1998; Tsukahara et al., 2010). Other CPC components, Borealin and Survivin, regulate SGO2 recruitment and pericentromeric localization (Tsukahara et al., 2010; Yamagishi et al., 2010). The catalytic component of the CPC is Aurora B kinase, which phosphorylates kinetochore substrates to destabilize microtubule interactions and is thus another potential drive effector. We find that positive selection shapes the domains of INCENP that interact with Borealin/Survivin, with HP1, and with Aurora B (Figure 5D), suggesting that INCENP can adapt to selfish centromere DNA by modulating its localization to pericentromeric heterochromatin and ultimately the recruitment of SGO2 and Aurora B.

Finally, in the effector module we find rapid evolution of SGO2, which is recruited by both the kinetochore and heterochromatin pathways (Figure 1B). This finding suggests that SGO2 can tune the relative strength of the two pathways through mutations that modulate its recruitment by either pathway. In comparison, SGO1 is a paralog of SGO2 that does not recruit MCAK (Yao and Dai, 2012) and does not have signatures of positive selection, suggesting that evolutionary pressure to regulate MCAK recruitment shapes SGO2 evolution. Overall, our molecular evolution analysis shows signatures of positive selection in both the kinetochore and heterochromatin pathways. We find these changes both in domains that interact directly with DNA and in protein-protein interaction domains leading to recruitment of drive effectors. These results are consistent with our parallel pathway model for drive and suppression, but not with a simpler model of an arms race limited to centromere DNA and DNA binding proteins.

## Discussion

Here we propose a parallel pathway model for drive and suppression of selfish centromeres: centromere DNA can exploit the kinetochore pathway to increase effector recruitment, and the host can make centromeres functionally equivalent by minimizing the contribution of the kinetochore pathway relative to the heterochromatin pathway (Figure 1F). This model predicts that disruption of either pathway will reduce effector (e.g., SGO2) recruitment, but the functional consequences will depend on which pathway is affected. Centromeres become either functionally more different if the symmetric heterochromatin pathway is weakened, or more similar if the asymmetric kinetochore pathway is weakened. In our experiments, either deletion of CENP-B or introduction of a divergent allele of CENP-C leads to SGO2 reduction to a similar extent (Figure 2A and Figure 3A). However, genetically different centromeres in CHPO hybrid oocytes become functionally more different when CENP-B is deleted (Figure 2E), whereas they become functionally more similar when rat CENP-C is expressed (Figure 3C). CENP-B deletion weakens the symmetric heterochromatin pathway, as shown by reduced H3K9me3, making the asymmetric kinetochore pathway more dominant. Loss of CENP-B also reduces CENP-C recruitment but does not affect the asymmetry between larger and smaller centromeres (Figure 2C). Complementing these findings, the CENP-C results are consistent with our model prediction that natural selection has acted on CENP-C interfaces involved in effector recruitment, so a divergent rat CENP-C interacts less well with mouse binding partners in the kinetochore pathway. Therefore, expression of rat CENP-C weakens the asymmetric kinetochore pathway, making the symmetric heterochromatin pathway relatively more dominant.

Our molecular evolution analysis shows adaptive evolution in multiple centromere proteins and in specific domains that interact with CENP-A chromatin or with other proteins leading to effector recruitment (Figure 4 and Figure 5). The previous model of a molecular arms race limited to interactions between centromere DNA and DNA-interacting proteins (such as CENP-A) (Henikoff et al., 2001) does not explain the more widespread recurrent evolution of centromere proteins. An alternative explanation, independent of centromere drive, is that the selective pressure may be related to non-centromere functions. For example, kinetochore proteins are repurposed for neural development in fly and worm (Cheerambathur et al., 2019; Zhao et al., 2019), and KNL1 (also known as CASC5) is implicated in human brain size regulation (Javed et al., 2018; Shi et al., 2016). However, such non-centromere functions have not been identified more broadly in eukaryotes. In contrast, our parallel pathway model predicts recurrent evolution of proteins in both pathways to equalize centromeres by weakening the kinetochore pathway or strengthening the heterochromatin pathway. In our model, selfish centromere DNA evolves to exploit the kinetochore pathway by recruiting more of a protein that ultimately recruits effectors. To suppress functional differences between centromeres, proteins in the kinetochore pathway can adapt to minimize the impact of selfish centromere DNA on kinetochore formation or effector recruitment. Furthermore, proteins in the heterochromatin pathway such as CENP-B and INCENP can adapt to increase effector recruitment equally at all centromeres, or SGO2 can adapt by modulating its recruitment by either pathway (Figure 1F). The acidic domain of CENP-B is implicated in recruiting heterochromatin proteins (Otake et al., 2020), and the number of negatively charged amino acids in this domain is recurrently changed in mammals (Supplementary Figure 2B and 2C). Although these changes are not analyzed in PAML, they suggest that CENP-B may have evolved to regulate pericentromeric heterochromatin. Overall, a protein network for effector recruitment can adapt to minimize asymmetric recruitment by selfish centromere DNA, while maintaining essential functions of the kinetochore and of microtubule destabilizing factors for accurate chromosome segregation.

Our results suggest an explanation for the conservation of CENP-B in mammals, as well as the presence of its binding sequence, the CENP-B box, at most mammalian centromeres with the notable exception of the Y chromosome. Although CENP-B is the only centromere protein known to bind a specific DNA sequence in mammals, neither the protein nor the binding sequence is essential for centromere function (Amor et al., 2004; Hudson et al., 1998; Kapoor et al., 1998; Logsdon et al., 2019; Perez-Castro et al., 1998). We propose that CENP-B is conserved because it suppresses functional differences between centromeres by strengthening the heterochromatin pathway (Figure 6), consistent with a more general function of heterochromatin in suppressing many selfish genetic elements (Allshire and Madhani, 2017). This CENP-B function is important only when centromeres of homologous chromosomes are different, which would frequently occur in outbred populations. Loss of CENP-B therefore increases functional difference between larger and smaller centromeres in our hybrid model, but does not significantly impair fertility or viability in inbred laboratory strains (Hudson et al., 1998; Kapoor et al., 1998; Perez-Castro et al., 1998). A potential cost of increasing heterochromatin, however, is that its invasion into CENP-A chromatin disrupts centromere function (Ohzeki et al., 2016). We therefore propose that mammalian CENP-B has acquired an additional function to maintain CENP-A chromatin, by recruiting CENP-C and CENP-A chromatin regulators (Fachinetti et al., 2015; Otake et al., 2020) (Figure 6). By regulating both CENP-A chromatin and heterochromatin, alternative functions of CENP-B in different chromatin environments may suppress functional differences between centromeres through heterochromatin while maintaining centromere function. CENP-B can suppress differences between centromeres only if its functions are insensitive to expansion of the number of CENP-B binding sites; otherwise it would contribute to higher levels of effector recruitment by DNA repeat expansions. Indeed, we find that CENP-B does not contribute to asymmetry in CENP-C recruitment between larger and smaller centromeres, despite 6-to 10-fold differences in minor satellite sequences containing CENP-B boxes (Iwata-Otsubo et al., 2017). This result suggests that CENP-B recruits CENP-C only within the CENP-A chromatin domain, so that CENP-B binding outside of this domain does not strengthen the kinetochore pathway (Figure 6). Furthermore, the heterochromatin symmetry between larger and smaller centromeres suggests that although CENP-B contributes to initiating heterochromatin formation, for example by recruiting an H3K9 methyltransferase, heterochromatin spreading does not depend on the number of CENP-B boxes (Figure 6). Initiation of heterochromatin propagation is a common mechanism to regulate heterochromatin formation, as in the example of X inactivation where XIST initiates heterochromatinization of the entire chromosome (Allshire and Madhani, 2017). Thus, CENP-B functions in CENP-A chromatin and heterochromatin are insensitive to repeat expansion. A centromere variant completely lacking CENP-B boxes, however, will lose to an existing centromere in female meiosis because it will recruit less effectors by both the kinetochore and heterochromatin pathways. Therefore, CENP-B boxes are maintained at most centromeres, but this selective pressure does not affect the Y chromosome, which never experiences female meiosis and does not bind CENP-B (Gamba and Fachinetti, 2020).

**Figure 6:**
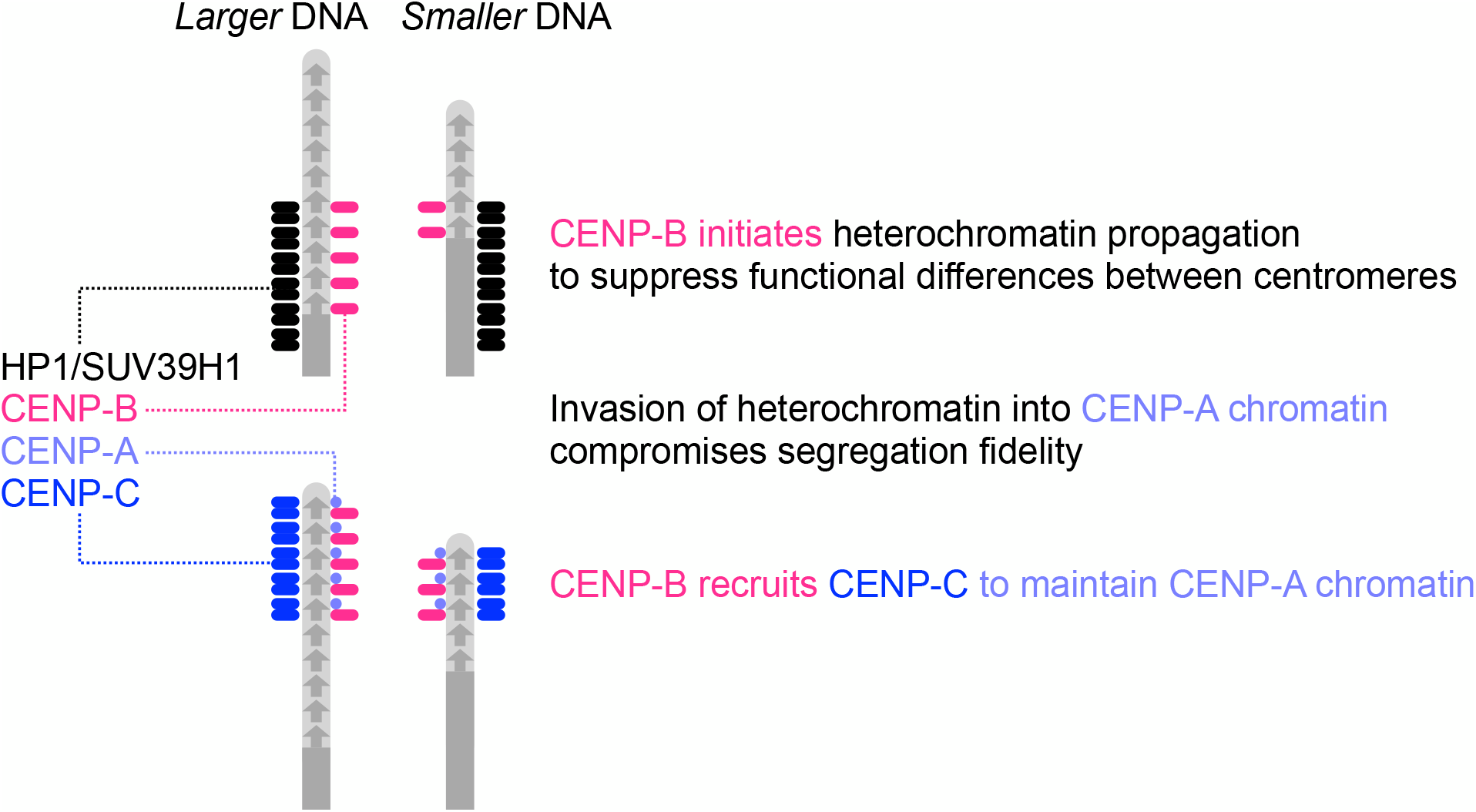
Dual functions of CENP-B in suppressing functional differences in centromeres while maintaining CENP-A chromatin. CENP-B initiates heterochromatin formation to equalize centromeres (top). Despite the difference in CENP-B binding sites, larger and smaller centromeres have similar amounts of H3K9me3 (Figure 1D), indicating that heterochromatin formation is insensitive to CENP-B abundance, likely due to self-propagation of heterochromatin. Invasion of heterochromatin into CENP-A compromises centromere function. To prevent this disruption, we propose that CENP-B has acquired an additional function in CENP-A chromatin (bottom): CENP-B recruits CENP-C but does not contribute to CENP-C asymmetry between larger and smaller centromeres (Figure 2C), suggesting that only CENP-B within CENP-A chromatin recruits CENP-C. Thus, CENP-B functions in heterochromatin and CENP-A chromatin are insensitive to repeat expansion.

Genetic conflict between selfish centromere DNA and centromere-binding proteins potentially explains the complexity of eukaryotic centromeres. Opportunities for selfish genetic elements to exploit the chromosome segregation machinery are not limited to female meiosis, as selfish plasmids (e.g., 2-micron plasmids in budding yeast) benefit by maximizing their transmission to daughter cells in mitosis (Malik and Henikoff, 2009; Rizvi et al., 2017). These opportunities are limited by the strong epigenetic component of most eukaryotic centromeres, which are not defined by specific DNA sequences. Centromeres cannot be completely independent of the underlying DNA sequence, however, because some protein must interact with DNA, so different sequences can have different binding affinities or impact the structure of the centromeric nucleosome complex (Allu et al., 2019). The presence of multiple pathways to form a kinetochore (e.g., via CENP-ACLN and CENP-TWSX connected by CENP-HIKM, or via CENP-OPQUR) (Cortes-Silva et al., 2020; Hamilton et al., 2020; Nishino et al., 2012; Pesenti et al., 2018; Veld et al., 2016; Weir et al., 2016; Yan et al., 2019) allows proteins to adapt by minimizing a pathway that is exploited by a selfish element, while maintaining kinetochore function via other pathways. Consistent with this idea of independent modules for kinetochore formation, CENP-A depletion leads to proportional reduction of centromeric CENP-C, whereas CENP-T and CENP-I persist longer (Fachinetti et al., 2013). In addition, recurrent changes in kinetochore modules are observed throughout eukaryotic evolution, such as changes in the number of MELT motifs in KNL1 and replacement of the SKA complex by the DAM complex (Hooff et al., 2017; Tromer et al., 2015). Regulation of kinetochore-microtubule attachment stability may be another way to suppress selfish genetic elements, as MELT motifs recruit BUB1 and SKA and DAM complexes stabilize attachments. Thus, internal conflicts between selfish genetic elements and the chromosome segregation machinery may have shaped complexity in eukaryotic centromeres.

## Acknowledgements

We thank B.E. Black for comments on the manuscript, Y. Watanabe for the CENP-C and SGO2 antibodies, J.B. Searle, C. Conroy (Museum of Vertebrate Zoology), and A. Ferguson (Field Museum of Natural History) for collecting Murinae tissue samples, and G. Thomas and J. Good for assistance with the genomic analyses. The research was supported by the NIH (R35GM122475 to M.A.L.; R35GM124684 to M.T.L.; RM1HG010023 to J.K.) and a predoctoral fellowship from the Funai Foundation for Information Technology (T.K.).

---

**Supplementary Figure 1:**
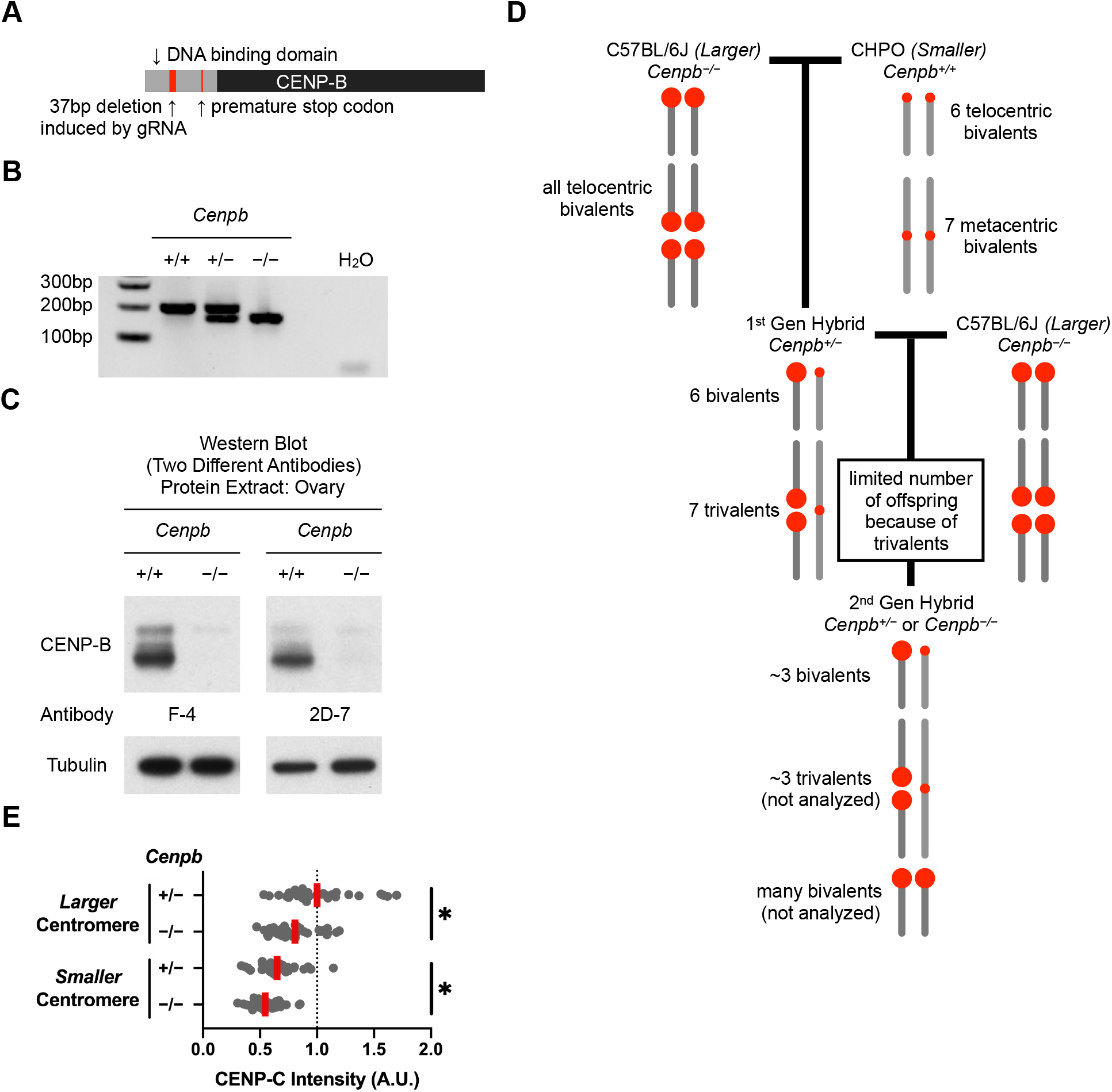
CRISPR genome editing creates CENP-B null mice. (A) Summary of CRISPR genome editing, using a gRNA designed to target the DNA binding domain of CENP-B. (B) CENP-B genotyping. As the CENP-B mutation is a 37bp deletion, a PCR reaction amplifying the flanking regions can distinguish three genotypes of CENP-B. (C) Absence of CENP-B protein in *Cenpb*^*−/−*^ mice. Protein extract from ovary is used to detect CENP-B using two different antibodies. (D) Detailed crossing scheme to produce second-generation hybrid *Cenpb*^*−/−*^ mice with larger and smaller paired centromeres (related to Figure 2B). The first cross produces first generation hybrid *Cenpb*^*+/−*^ animals with smaller centromeres inherited from CHPO. Because CHPO has six telocentric chromosomes and seven metacentrics formed by Robertsonian chromosome fusions, the first-generation hybrids contain six bivalents in meiosis and seven trivalents, in which a Robertsonian fusion from CHPO pairs with two homologous telocentric chromosomes (Chmátal et al., 2014). Trivalents are associated with meiotic errors (Bint et al., 2011; Daniel, 2002; Pacchierotti et al., 1995), and the first-generation hybrids exhibit low fertility, but some progeny can be obtained in a second cross to *Cenpb*^*−/−*^. These second-generation hybrids inherit some smaller centromeres from the first-generation hybrid parent, and 25% are *Cenpb*^*−/−*^ females that can be used to collect oocytes for our analyses. Oocytes from the second-generation hybrids do not arrest at metaphase I, likely because they have fewer trivalents that activate the spindle assembly checkpoint (Chmátal et al., 2015). Therefore, we are unable to measure biased orientation of larger centromeres towards the egg side of the spindle, as previously reported in first-generation hybrids (Iwata-Otsubo et al., 2017), because this bias depends on delayed progression through meiosis I (Akera et al., 2019). (E) CENP-C reduction in the second generation hybrid (related to Figure 2C). Oocytes from the second generation hybrid (Figure 2B) were microinjected with cRNA for GFP-tagged dCas9 and gRNA targeting minor satellite centromere DNA, fixed at metaphase I, and stained for CENP-C. Each dot represents a single centromere (n=34 centromeres for each construct); red line, mean; *p<0.01.

**Supplementary Figure 2:**
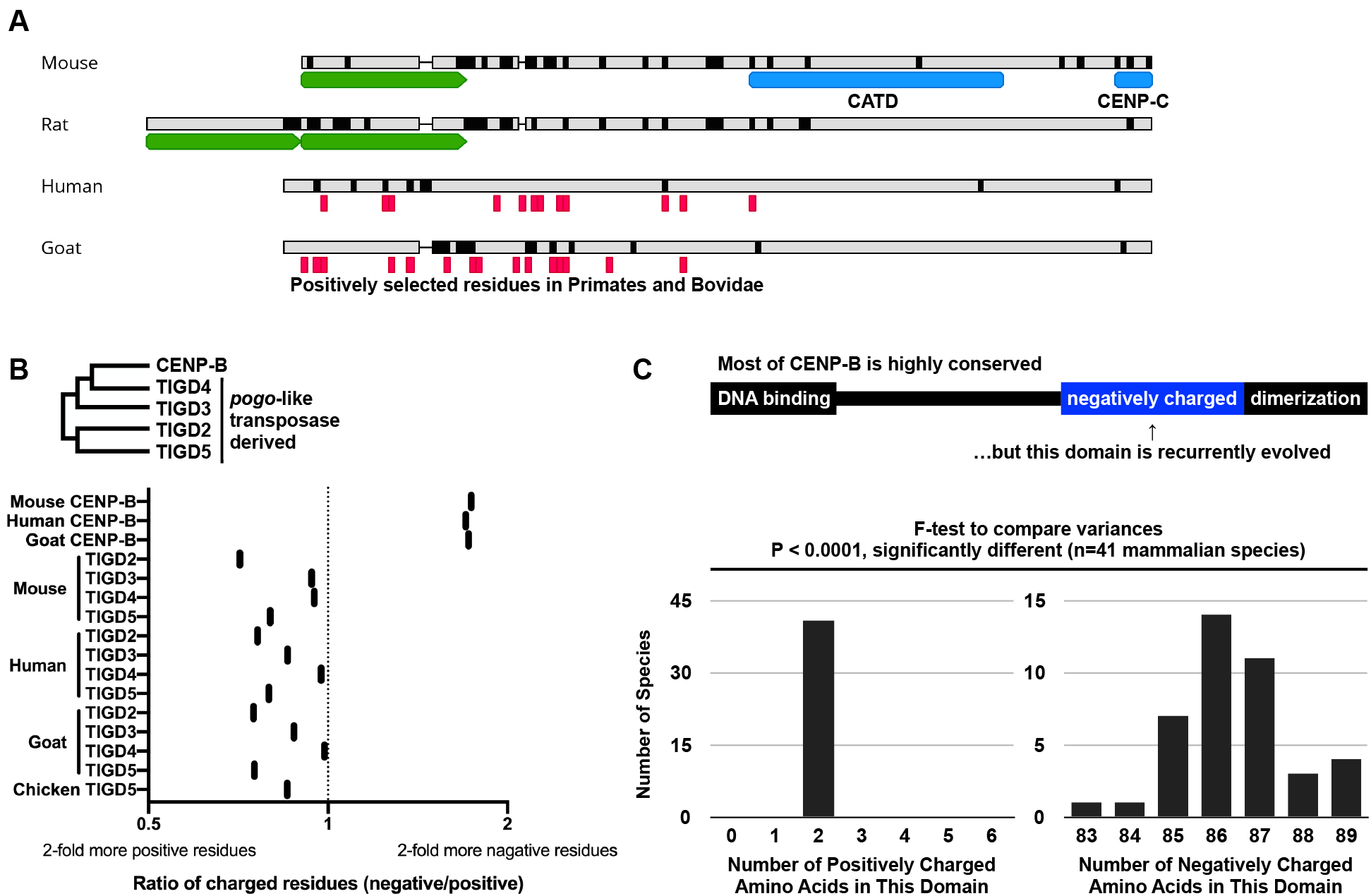
Changes in CENP-A and CENP-B cannot be analyzed by standard method to detect adaptive evolution. (A) Changes in CENP-A N-terminal tails. CENP-A amino acid sequences of four mammalian species are aligned. Known domains of CENP-A are shown in blue boxes, deviation from the consensus sequence of all four species is shown in black, and deletions are shown as thin lines. Signatures of positive selection were previously found in primate CENP-A (Schueler et al., 2010), shown in red boxes in the human sequence. Bovine genomes (Chen et al., 2019) are used to detect signatures of positive selection in CENP-A, and the result is shown in the goat sequence. Such signatures are mostly found in the N-terminal tail. The N-terminal tail of Murinae CENP-A is either short (as in mouse) or long with two tandem duplicates (as in rat) (green boxes). Thus, alignment of the Murinae CENP-A N-terminal tail is difficult and removed from our PAML analysis. (B) CENP-B negatively charged domain. Mouse, human, and goat are shown as examples of genomes with CENP-B and paralogous pogo-like transposases. The ratio of negatively charged to positively charged amino acids is plotted. As pogo-like transposases have fewer negatively charged amino acids than CENP-B, the negatively charged domain is likely unique to CENP-B. (C) Changes in the CENP-B negatively charged domain. Although most of CENP-B is highly conserved, the number of negatively charged amino acids is variable in mammals. For comparison, the number of positively charged amino acids does not change in this domain. The number of species for each number of positively charged or negatively charged amino acids in this domain is plotted.

**Supplementary Table 1.**

| Genome | *Ha* | *Pd* | *Mn* | *Gd* | *Rd* | *Rs* |
| --- | --- | --- | --- | --- | --- | --- |
| LongRanger reference-guided assembly |  |  |  |  |  |  |
| Mean Molecule Length (kb)<br>[Input DNA Quality] | 8.7 | 11.3 | 16.7 | 8.9 | 13.8 | 16.8 |
| DNA in Molecules >100kb (%)<br>[Input DNA Quality] | 20.9 | 19.4 | 24.0 | 14.7 | 10.1 | 12.0 |
| Mean DNA per GEM (kb)<br>[Linked-Reads Quality] | 288.1 | 638.0 | 593.5 | 447.5 | 739.4 | 654.7 |
| N50 Linked-Reads per Molecule<br>[Linked-Reads Quality] | 5.0 | 4.0 | 4.0 | 5.0 | 6.0 | 9.0 |
| N50 Phase Block (kb)<br>[Assembly Quality] | 10.2 | 28.2 | 28.5 | 42.9 | 37.6 | 115.4 |
| Number of Reads (million reads) | 1378 | 1518 | 1389 | 1413 | 1415 | 1444 |
| Mean Depth | 37.7 | 44.6 | 38.9 | 33.9 | 34.3 | 39.3 |
| Mapped Reads (%) | 69.1 | 72.3 | 68.8 | 63.0 | 65.3 | 69.3 |
| Supernova *de novo* assembly |  |  |  |  |  |  |
| Number of Reads (million reads) | 1378 | 800 | 1389 | 800 | 1415 | 1444 |
| Number of Scaffolds ≥10 kb (K) | 25.23 | 66.72 | 71.61 | 28.98 | 8.72 | 2.54 |
| N50 Contig Size (kb) | 13.31 | 19.47 | 17.16 | 12.19 | 28.42 | 82.08 |
| N50 Phase Block Size (kb) | 4.19 | 29.01 | 25.74 | 7.65 | 202.77 | 880.7 |
| N50 Scaffold Size (Mb) | 15<br>kb | 31<br>kb | 25<br>kb | 14<br>kb | 2.67 | 8.73 |
| Assembly Size<br>(Only Scaffolds ≥10 kb) (Gb) | 393<br>Mb | 1.72 | 1.60 | 410<br>Mb | 2.29 | 2.26 |
| Effective Coverage | 31.2 | 26.3 | 33.2 | 26.3 | 55.8 | 56.4 |
| Estimated Genome Size (Gb) | 3.6 | 3.3 | 4.0 | 3.2 | 2.7 | 2.7 |

**Supplementary Table 2.**
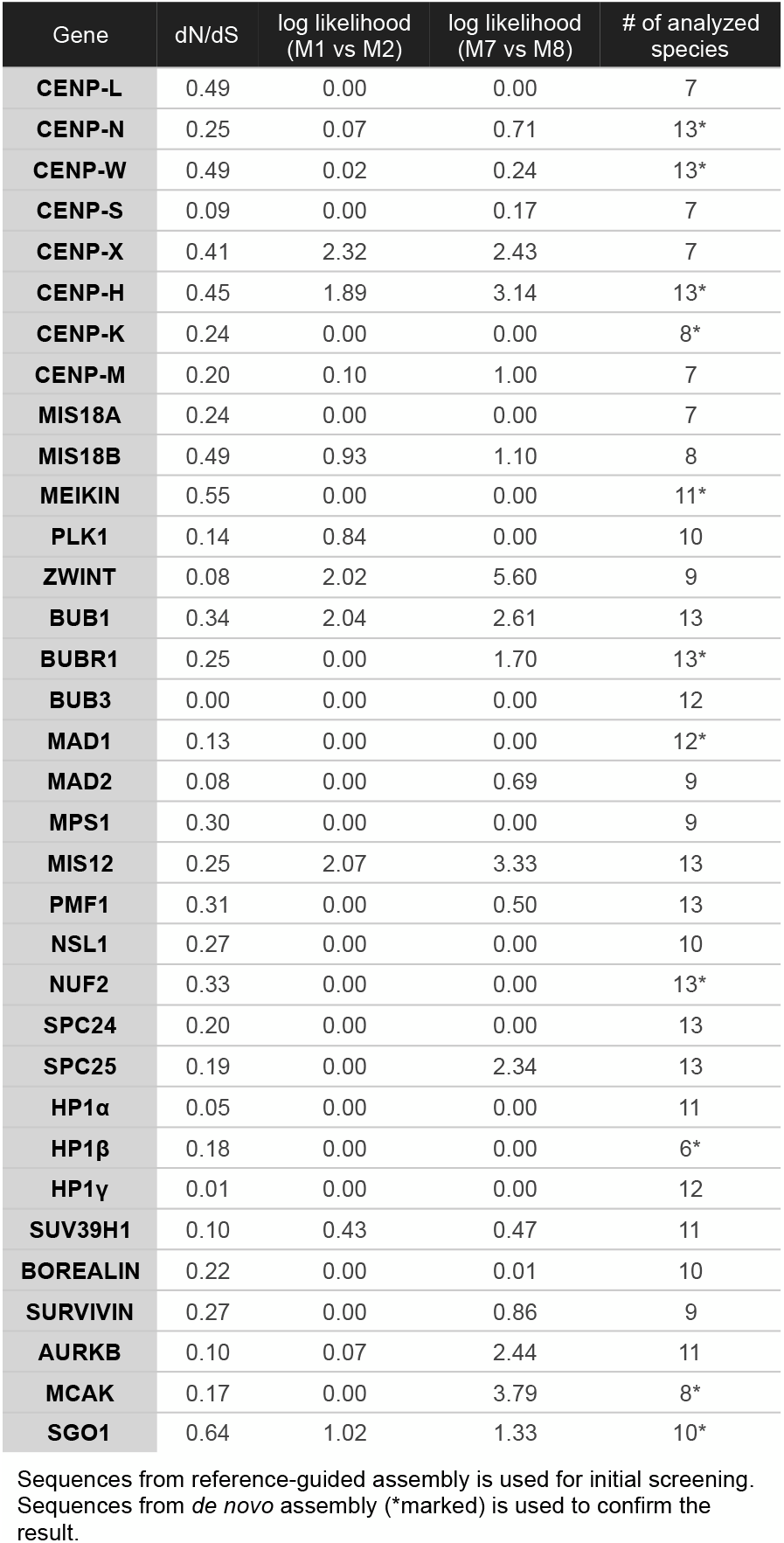

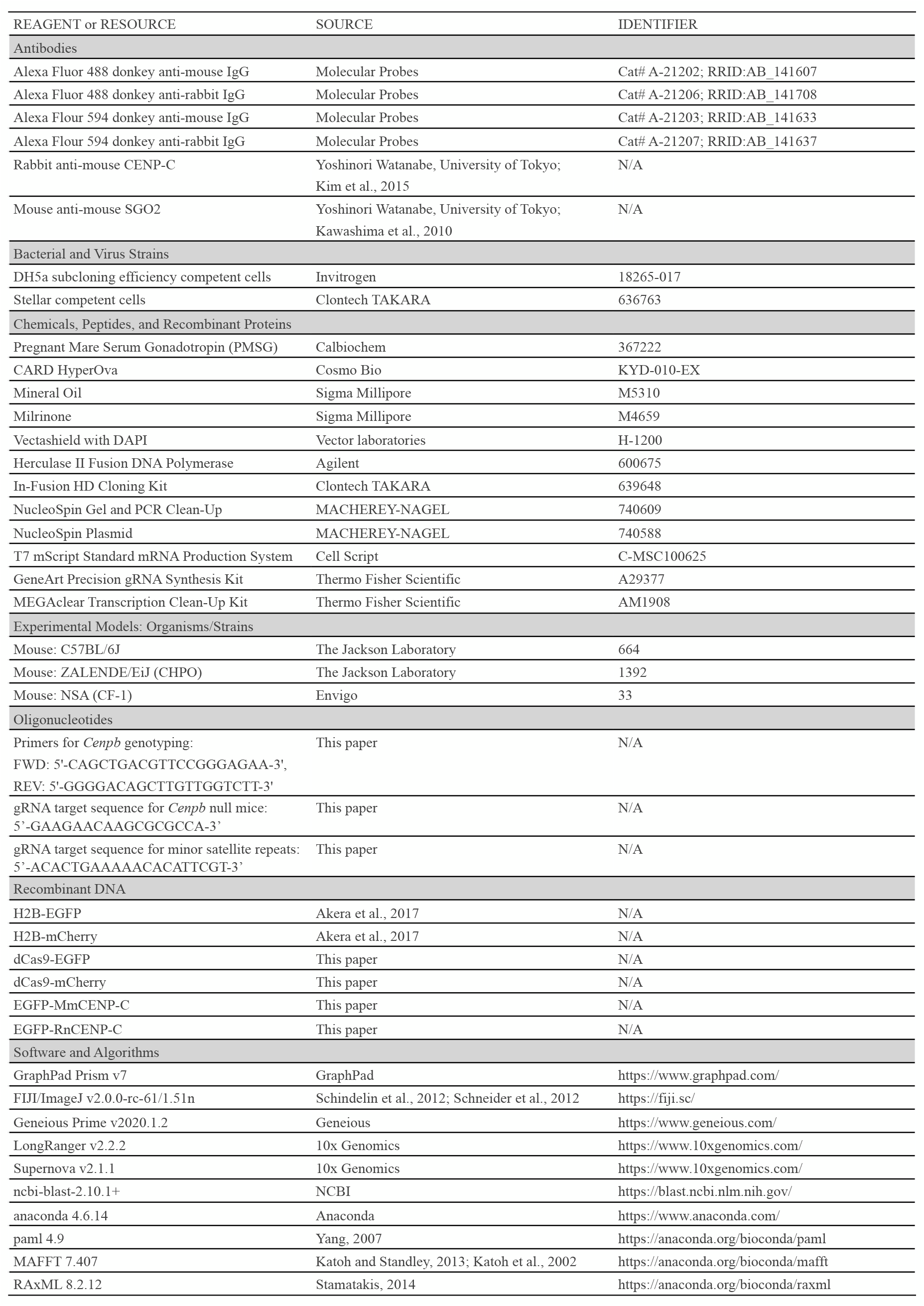
KEY RESOURCES TABLE

## LEAD CONTACT AND MATERIALS AVAILABILITY

Further information and requests for resources and reagents should be directed to and will be fulfilled by the Lead Contact, Michael A. Lampson.

## EXPERIMENTAL MODEL AND SUBJECT DETAILS

### Mice

Mouse strains were purchased from the Jackson Laboratory (ZALENDE/EiJ, stock #001392 corresponds to CHPO; C57BL/6J, stock# 000664) and from Envigo (NSA, stock# 033 corresponds to CF-1). CHPO males were crossed to CF-1 females to generate hybrids shown in Figure 1c. The CHPO strain contains seven Robertsonian fusions (Rb(1.3), Rb(4.6), Rb(5.15), Rb(11.13), Rb(8.12), Rb(9.14), and Rb(16.17)), each of which pairs with two CF-1 chromosomes in CHPO hybrid meiosis I to form a trivalent (Chmátal et al., 2014). We included only bivalents (chromosome 2, 7, 10, 18, 19, X) in our analyses to avoid complications of trivalents.

In order to generate CENP-B null mice, 1-cell embryos (from female CF-1 and male DBA/2J x C57BL/6J hybrid) were collected and microinjected with Cas9 mRNA (TriLink, CleanCap Cas9 mRNA, L-7606) and gRNA (GAAGAACAAGCGCGCCA) (Thermo Fisher scientific, GeneArt Precision gRNA Synthesis Kit, A29377). Embryos were cultured *in vitro* until blastocyst stage and transferred to pseudopregnant females to produce a founder mouse carrying 37bp deletion (TGAGCACCATCCTGAAGAACAAGCGCGCCATCCTGGC) that produces a premature stop codon at Leu100 in the DNA binding domain. The founder was crossed with C57BL/6J for multiple generations to remove possible off-target mutations. Mice were genotyped by extracting genomic DNA from tail clip (QIAGEN, DNeasy Blood & Tissue Kit, 69504) and amplifying a *Cenpb* fragment (Agilent, Herculase II Fusion DNA Polymerase). To generate *Cenpb* null mice with larger and smaller centromeres, CHPO females were crossed to C57BL/6J *Cenpb* null males to generate first generation hybrid females, which were then crossed to C57BL/6J *Cenpb* null males to generate second generation hybrid females as shown in Figure 2B and Supplementary Figure 1D. All animal experiments were approved by the Institutional Animal Care and Use Committee and were consistent with the National Institutes of Health guidelines.

## METHOD DETAILS

### Oocyte collection and culture

Female mice (8-14 weeks of age) were hormonally primed with 5U of Pregnant Mare Serum Gonadotropin (PMSG, Calbiochem, cat# 367222) or 0.1mL of CARD HyperOva (Cosmo Bio, KYD-010-EX) 44-48 h prior to oocyte collection. Germinal vesicle (GV)-intact oocytes were collected in M2 medium (Sigma, M7167), denuded from cumulus cells, and cultured in Chatot-Ziomek-Bavister (CZB) medium (Thermo Fisher, MR019D) in a humidified atmosphere of 5% CO2 in air at 37.8C°. During collection, meiotic resumption was inhibited by addition of 2.5 mM milrinone. Milrinone was subsequently washed out to allow meiotic resumption. Oocytes were checked for GVBD (germinal vesicle breakdown), and those that did not enter GVBD stage were removed from the culture.

### Oocyte microinjection

GV oocytes were microinjected with ~5 pl of cRNAs in M2 medium (with 2.5 mM milrinone and 3mg/mL BSA) at room temperature (RT) with a micromanipulator TransferMan NK 2 (Eppendorf) and picoinjector (Medical Systems Corp.). After the injection, oocytes were kept in milrinone for 16 h to allow protein expression. cRNAs used for microinjections were dCas9-EGFP (dead Cas9 with EGFP at the N terminus) at 1000ng/μL, dCas9-mCherry (dead Cas9 with mCherry at the N terminus) at 1000ng/μL, gRNA that targets minor satellite repeat (ACACTGAAAAACACATTCGT) at 200ng/μL, H2B-EGFP (human histone H2B with EGFP at the C terminus) at 150ng/μL, H2B-mCherry (human histone H2B with mCherry at the C terminus) at 150ng/μL, EGFP-MmCENP-C (mouse CENP-C with EGFP at the N terminus) at 100ng/μL, and EGFP-RnCENP-C (rat CENP-C with EGFP at the N terminus) at 100ng/μL. cRNAs were synthesized using the T7 mScriptTM Standard mRNA Production System (CELL SCRIPT) or mMESSAGE mMACHINE SP6 Transcription Kit (Thermo Fisher scientific). gRNAs were synthesized using GeneArt Precision gRNA Synthesis Kit (Thermo Fisher scientific A29377).

### Live imaging and chromosome position assay

For the chromosome position assay, oocytes were collected and microinjected with the constructs indicated in the figure legends. After inducing meiotic resumption by washing out milrinone, oocytes were placed into 2μL drops of CZB media covered with mineral oil in a glass-bottom tissue culture dish (FluoroDish FD35-100) in a heated environmental chamber with a stage top incubator (Incubator BL and Heating Insert P; PeCon GmBH) to maintain 37C°. Confocal images were collected with a microscope (DMI4000 B; Leica) equipped with a 63x 1.3 NA glycerol-immersion objective lens, an xy piezo Z stage (Applied Scientific Instrumentation), a spinning disk confocal scanner (Yokogawa Corporation of America), an electron multiplier charge-coupled device camera (ImageEM C9100-13; Hamamatsu Photonics), and an LMM5 laser merge module with 488- and 593-nm diode lasers (Spectral Applied Research) controlled by MetaMorph software (Molecular Devices). Confocal images were collected as z stacks at 0.5 μm intervals to visualize the entire meiotic spindle. The position of the spindle near the cortex was confirmed by differential interference contrast images. The spindle equator was determined as a middle of the spindle. The chromosome position of each bivalent was determined as a crossover site and normalized by the distance between spindle equator and spindle poles.

### Oocyte immunocytochemistry

After inducing meiotic resumption by washing out milrinone (4.5 hours for prometaphase staining and 7.5 hours for metaphase staining), MI oocytes were fixed in freshly prepared 2% paraformaldehyde in PBS with 0.1% Triton X-100, pH 7.4, for 20 min at RT, permeabilized in PBS with 0.1% Triton X-100 for 15 min at RT, placed in blocking solution (PBS containing 0.3% BSA and 0.01% Tween-20) 15 min RT or overnight at 4C, incubated 1-2 h with primary antibodies in blocking solution, washed 3 times for 15 min each, incubated 1 h with secondary antibodies, washed 3 times for 15 min each, and mounted in Vectashield with DAPI (Vector, H-1200) to visualize chromosomes. Primary antibodies used for this study were rabbit anti-human H3K9me3 (1:500; Abcam, ab8898), mouse anti-mouse SGO2 (1:500, a gift from Yoshinori Watanabe), and rabbit anti-mouse CENP-C (1:2500, a gift from Yoshinori Watanabe). Secondary antibodies were Alexa Fluor 488–conjugated donkey anti-rabbit or donkey anti-mouse or Alexa Fluor 594–conjugated donkey anti-rabbit, or donkey anti-mouse (1:500, Invitrogen). Confocal images were collected as z stacks at 0.5 μm intervals to visualize the entire meiotic spindle, using the spinning disc confocal microscope described above. To quantify centromere signal ratios, optical slices containing centromeres from the same bivalent were added to produce a sum projection using Fiji/ImageJ. Ellipses were drawn around the centromeres, and signal intensity was integrated over each ellipse after subtracting cytoplasmic background. Ratios were obtained for each bivalent by dividing the intensity of the larger centromere by that of the smaller centromere, as determined by dCas9 signal intensity.

### Whole Genome Sequencing of Six Murinae Species

Frozen tissue samples from male individuals were obtained from the Museum of Vertebrate Zoology, Berkeley, CA (MZV) and the Field Museum of Natural History, Chicago, IL (FMNH). *Hylomyscus alleni* (MVZ Mamm 196246) was captured in Cameroon in 2000, *Praomys delectorum* (MVZ Mamm 221157) was captured in Malawi in 2007, *Mastomys natalensis* (MVZ Mamm 221054) was captured in Malawi in 2007, *Grammomys dolichurus* (MVZ Mamm 221001) was captured in Malawi in 2007, *Rhabdomys dilectus* (FMNH 192475) was captured in Malawi in 2006, and *Rhynchomys soricoides* (FMNH 198792) was captured in The Philippines in 2008. All genomes were sequenced in the Center for Applied Genomics at Children’s Hospital of Philadelphia. High molecular weight DNA was extracted following the protocol provided by 10xGenomics (CG000072 Rev B Sample Preparation Demonstrated Protocol, DNA Extraction from Fresh Frozen Tissue). Extracted DNA was quality controlled (CG00019 Rev B Sample Preparation Demonstrated Protocol, High Molecular Weight DNA QC), and all of the samples had a mean length greater than 50kb, and high enough concentration to dilute to 1ng/μL for library preparation. Chromium Genome Reagent Kits v2 from 10xGenomics was used to prepare libraries of 2×150 base reads, with read 1 constituting 10xBarcode (16bp) + nmer (6bp) + genome sequence (128bp) and read 2 constituting genome sequence (150bp). i7 index used 8bp sample index, and i5 index was not used. Sequencing depth was calculated based on putative genome size 3Gb and coverage 56x, following 10xGenomics R&D recommendation, and the libraries were sequenced with Illumina HiSeq. Demultiplexed FASTQ files were analyzed using the LongRanger wgs - basic pipeline. This pipeline gave general QC statistics related to the 10x barcoding and number of read pairs present in the FASTQ files. All sample FASTQs contained more than 688M read pairs and have acceptable barcode diversity/% on whitelist. LongRanger was used to assemble genomes, using the *Mus musculus* (mm10) as reference. In parallel, Supernova was used to assemble *de novo* genomes. See Supplementary Table 1 for assembly statistics. In order to obtain protein coding sequences, mm10 annotation was used to annotate reference-guided assemblies, and translated BLAST (tblastn) was used to pull homologous sequences from de novo assemblies using *Mus musculus* protein sequences as query sequences.

### Phylogenetic Tree Construction

The species tree shown in Figure 4B was obtained from maximum likelihood (RAxML) and Bayesian inference (MrBayes). The phylogeny within *Mus* was previously studied (Keane et al., 2011; Thybert et al., 2018). In order to resolve phylogeny in Murinae, the same set of genes that were used to construct a primate phylogenetic tree (Perelman et al., 2011) was aligned by MAFFT (Katoh and Standley, 2013; Katoh et al., 2002). The initial alignment was imported in Geneious Prime, and manually inspected for sequence alignment ambiguity. Ambiguous regions were removed from subsequent analyses. Maximum likelihood tree was constructed with RAxML (Stamatakis, 2014), and Bayesian inference tree was constructed with MrBayes (Huelsenbeck and Ronquist, 2001), with *Peromyscus maniculatus* as outgroup. Both inferences supported the tree topology shown in Figure 4B.

### Molecular Evolution Analyses

In order to create a histogram in Figure 4C, alignments of mouse-rat orthologs were filtered for dS below 0.5, as higher dS values indicate misalignment. A list of genes for each subcellular compartment was obtained from Human Protein Atlas. Mouse-human orthologs were used to calculate average dN/dS for each subcellular compartment in Figure 4D. The analysis to identify signatures of positive selection (PAML) is highly sensitive to alignment errors, so automated genome-wide analysis is prone to false positives (van der Lee et al., 2017). To prevent these errors, alignments for selected genes were manually inspected. Coding sequences for each gene were aligned by Geneious Alignment (translation align) implemented in Geneious Prime, and manually inspected for sequence alignment ambiguity. Insertions or deletions as well as their flanking codons were removed from analyses. To test signatures of positive selection, we compared the likelihood of models of neutral codon evolution to models of codon evolution allowing positive selection, implemented in PAML version 4 (Yang, 2007). The neutral model M1 (fixed dN/dS values between 0 to 1) and M2 (M1 parameters plus dN/dS > 1) were compared in the first test, and the neutral model M7 (dN/dS values fit a beta distribution from 0 to 1) and M8 (M7 parameters plus dN/dS > 1) were compared in the second test, assuming the F3×4 model of codon frequencies. Degree of freedom for each test was 2, and the log likelihood test was significant above 5.99 (p < 0.05). We first used the species tree, and signatures of positive selection were confirmed using a gene tree for each gene, created by RAxML.

## QUANTIFICATION AND STATISTICAL ANALYSIS

Data points are pooled from at least two independent experiments. The following statistical methods were used: unpaired t test in Figures 2A, 2C, 2E, 3A, 3B, 3C, and S1E; Mann-Whitney U test in Figure 4D; chi square test for goodness of fit for deviations from 1 in Figure 1D and for statistical models (likelihood-ratio test) in Figure 4E and Supplementary Table 2; Naïve Emprical Bayes (NEB) analysis and Bayes Empirical Bayes (BEB) analysis in Figures 4B and 4E; F test to compare variance in Figure S2C. The exact value of n, what n represents, and definition of center can be found in the figure legends for each experiment. Unpaired t test, Mann-Whitney U test, and F test were performed using GraphPad Prism; chi square tests were performed using Excel; NEB and BEB analyses were performed using PAML model 2 and 8. *P* value of less than 0.05 was judged as statistically significant.

## DATA AND CODE AVAILABILITY

The draft genomes and raw sequencing reads have been submitted to the NCBI BioProject database (https://www.ncbi.nlm.nih.gov/bioproject) under accession number PRJNA669840. In-house scripts and pipelines are available from the authors upon request. Raw imaging data is available from the authors upon request.

## AUTHOR CONTRIBUTIONS

Conceptualization, T.K., M.T.L. and M.A.L.; Methodology, T.K.; Software, T.K., D.S. and E.C.N.; Investigation, T.K., J.M. and R.B.A.; Writing – Original Draft, T.K.; Writing – Review & Editing, T.K., M.T.L. and M.A.L.; Funding Acquisition, T.K., J.K., M.L.T. and M.A.L.; Resources, J.K. and M.A.L; Supervision, M.A.L.

## References

Abe, Y., Sako, K., Takagaki, K., Hirayama, Y., Uchida, K.S.K., Herman, J.A., DeLuca, J.G., and Hirota, T. (2016). HP1-Assisted Aurora B Kinase Activity Prevents Chromosome Segregation Errors. Dev Cell 36, 487–497.

Ainsztein, A.M., Kandels-Lewis, S.E., Mackay, A.M., and Earnshaw, W.C. (1998). INCENP Centromere and Spindle Targeting: Identification of Essential Conserved Motifs and Involvement of Heterochromatin Protein HP1. J Cell Biol 143, 1763–1774.

Akera, T., Chmátal, L., Trimm, E., Yang, K., Aonbangkhen, C., Chenoweth, D.M., Janke, C., Schultz, R.M., and Lampson, M.A. (2017). Spindle asymmetry drives non-Mendelian chromosome segregation. Science 358, 668–672.

Akera, T., Trimm, E., and Lampson, M.A. (2019). Molecular Strategies of Meiotic Cheating by Selfish Centromeres. Cell 178, 1132–1144.e10.

Allshire, R.C., and Madhani, H.D. (2017). Ten principles of heterochromatin formation and function. Nat Rev Mol Cell Biology 19, 229–244.

Allu, P.K., Dawicki-McKenna, J.M., Eeuwen, T.V., Slavin, M., Braitbard, M., Xu, C., Kalisman, N., Murakami, K., and Black, B.E. (2019). Structure of the Human Core Centromeric Nucleosome Complex. Curr Biol 29, 2625–2639.e5.

Amor, D.J., Bentley, K., Ryan, J., Perry, J., Wong, L., Slater, H., and Choo, K.H.A. (2004). Human centromere repositioning “in progress”. Proc National Acad Sci 101, 6542–6547.

Bint, S.M., Ogilvie, C.M., Flinter, F.A., Khalaf, Y., and Scriven, P.N. (2011). Meiotic segregation of Robertsonian translocations ascertained in cleavage-stage embryos--implications for preimplantation genetic diagnosis. Hum Reprod 26, 1575–1584.

Casola, C., Hucks, D., and Feschotte, C. (2007). Convergent Domestication of pogo-like Transposases into Centromere-Binding Proteins in Fission Yeast and Mammals. Mol Biol Evol 25, 29–41.

Cheerambathur, D.K., Prevo, B., Chow, T.-L., Hattersley, N., Wang, S., Zhao, Z., Kim, T., Gerson-Gurwitz, A., Oegema, K., Green, R., et al. (2019). The Kinetochore-Microtubule Coupling Machinery Is Repurposed in Sensory Nervous System Morphogenesis. Dev Cell 48, 864–72.e7.

Chen, L., Qiu, Q., Jiang, Y., Wang, K., Lin, Z., Li, Z., Bibi, F., Yang, Y., Wang, J., Nie, W., et al. (2019). Large-scale ruminant genome sequencing provides insights into their evolution and distinct traits. Science 364, eaav6202.

Chmátal, L., Gabriel, S.I., Mitsainas, G.P., Martínez-Vargas, J., Ventura, J., Searle, J.B., Schultz, R.M., and Lampson, M.A. (2014). Centromere strength provides the cell biological basis for meiotic drive and karyotype evolution in mice. Curr Biology Cb 24, 2295–2300.

Chmátal, L., Yang, K., Schultz, R.M., and Lampson, M.A. (2015). Spatial Regulation of Kinetochore Microtubule Attachments by Destabilization at Spindle Poles in Meiosis I. Curr Biology Cb 25, 1835–1841.

Cortes-Silva, N., Ulmer, J., Kiuchi, T., Hsieh, E., Cornilleau, G., Ladid, I., Dingli, F., Loew, D., Katsuma, S., and Drinnenberg, I.A. (2020). CenH3-Independent Kinetochore Assembly in Lepidoptera Requires CCAN, Including CENP-T. Curr Biology Cb 30, 561–572.e10.

Dambacher, S., Deng, W., Hahn, M., Sadic, D., Fröhlich, J., Nuber, A., Hoischen, C., Diekmann, S., Leonhardt, H., and Schotta, G. (2012). CENP-C facilitates the recruitment of M18BP1 to centromeric chromatin. Nucleus 3, 101–110.

Daniel, A. (2002). Distortion of female meiotic segregation and reduced male fertility in human Robertsonian translocations: Consistent with the centromere model of co-evolving centromere DNA/centromeric histone (CENP-A). Am J Med Genet 111, 450–452.

Dunleavy, E.M., Roche, D., Tagami, H., Lacoste, N., Ray-Gallet, D., Nakamura, Y., Daigo, Y., Nakatani, Y., and Almouzni-Pettinotti, G. (2009). HJURP Is a Cell-Cycle-Dependent Maintenance and Deposition Factor of CENP-A at Centromeres. Cell 137, 485–497.

Echave, J., Spielman, S.J., and Wilke, C.O. (2016). Causes of evolutionary rate variation among protein sites. Nat Rev Genetics 17, 109–121.

Fachinetti, D., Folco, H.D., Nechemia-Arbely, Y., Valente, L.P., Nguyen, K., Wong, A.J., Zhu, Q., Holland, A.J., Desai, A., Jansen, L.E.T., et al. (2013). A two-step mechanism for epigenetic specification of centromere identity and function. Nat Cell Biol 15, 1056–1066.

Fachinetti, D., Han, J.S., McMahon, M.A., Ly, P., Abdullah, A., Wong, A.J., and Cleveland, D.W. (2015). DNA Sequence-Specific Binding of CENP-B Enhances the Fidelity of Human Centromere Function. Dev Cell 33, 314–327.

Finseth, F.R., Dong, Y., Saunders, A., and Fishman, L. (2015). Duplication and Adaptive Evolution of a Key Centromeric Protein in Mimulus, a Genus with Female Meiotic Drive. Mol Biol Evol 32, 2694–2706.

Finseth, F.R., Nelson, T.C., and Fishman, L. (2020). Selfish chromosomal drive shapes recent centromeric histone evolution in monkeyflowers.

Fishman, L., and Saunders, A. (2008). Centromere-associated female meiotic drive entails male fitness costs in monkeyflowers. Sci New York N Y 322, 1559–1562.

Foltz, D.R., Jansen, L.E.T., Bailey, A.O., Yates, J.R., Bassett, E.A., Wood, S., Black, B.E., and Cleveland, D.W. (2009). Centromere-Specific Assembly of CENP-A Nucleosomes Is Mediated by HJURP. Cell 137, 472–484.

Fujita, Y., Hayashi, T., Kiyomitsu, T., Toyoda, Y., Kokubu, A., Obuse, C., and Yanagida, M. (2007). Priming of Centromere for CENP-A Recruitment by Human hMis18α, hMis18β, and M18BP1. Dev Cell 12, 17–30.

Gamba, R., and Fachinetti, D. (2020). From evolution to function: Two sides of the same CENP-B coin? Exp Cell Res 390, 111959.

Gao, B., Wang, Y., Diaby, M., Zong, W., Shen, D., Wang, S., Chen, C., Wang, X., and Song, C. (2020). Evolution of pogo, a separate superfamily of IS630-Tc1-mariner transposons, revealing recurrent domestication events in vertebrates. Mobile Dna-Uk 11, 25.

Gibbs, R.A., Weinstock, G.M., Metzker, M.L., Muzny, D.M., Sodergren, E.J., Scherer, S., Scott, G., Steffen, D., Worley, K.C., Burch, P.E., et al. (2004). Genome sequence of the Brown Norway rat yields insights into mammalian evolution. Nature 428, 493–521.

Godek, K.M., Kabeche, L., and Compton, D.A. (2014). Regulation of kinetochore-microtubule attachments through homeostatic control during mitosis. Nat Rev Mol Cell Biology 16, 57–64.

Hamilton, G.E., Helgeson, L.A., Noland, C.L., Asbury, C.L., Dimitrova, Y.N., and Davis, T.N. (2020). Reconstitution reveals two paths of force transmission through the kinetochore. Elife 9, e56582.

Henikoff, S., Ahmad, K., and Malik, H.S. (2001). The Centromere Paradox: Stable Inheritance with Rapidly Evolving DNA. Science 293, 1098–1102.

Hooff, J.J. van, Snel, B., and Kops, G.J. (2017). Unique phylogenetic distributions of the Ska and Dam1 complexes support functional analogy and suggest multiple parallel displacements of Ska by Dam1. Genome Biol Evol 9, 1295–1303.

Hudson, D.F., Fowler, K.J., Earle, E., Saffery, R., Kalitsis, P., Trowell, H., Hill, J., Wreford, N.G., Kretser, D.M. de, Cancilla, M.R., et al. (1998). Centromere Protein B Null Mice are Mitotically and Meiotically Normal but Have Lower Body and Testis Weights. J Cell Biol 141, 309–319.

Huelsenbeck, J.P., and Ronquist, F. (2001). MRBAYES: Bayesian inference of phylogenetic trees. Bioinformatics 17, 754–755.

Iwata-Otsubo, A., Dawicki-McKenna, J.M., Akera, T., Falk, S.J., Chmátal, L., Yang, K., Sullivan, B.A., Schultz, R.M., Lampson, M.A., and Black, B.E. (2017). Expanded Satellite Repeats Amplify a Discrete CENP-A Nucleosome Assembly Site on Chromosomes that Drive in Female Meiosis. Curr Biology Cb 27, 2365–2373.e8.

Jangam, D., Feschotte, C., and Betrán, E. (2017). Transposable Element Domestication As an Adaptation to Evolutionary Conflicts. Trends Genet 33, 817–831.

Javed, A.O., Li, Y., Muffat, J., Su, K.-C., Cohen, M.A., Lungjangwa, T., Aubourg, P., Cheeseman, I.M., and Jaenisch, R. (2018). Microcephaly Modeling of Kinetochore Mutation Reveals a Brain-Specific Phenotype. Cell Reports 25, 368–382.e5.

Kapoor, M., Luna, R.M. de O., Liu, G., Lozano, G., Cummings, C., Mancini, M., Ouspenski, I., Brinkley, B.R., and May, G.S. (1998). The cenpB gene is not essential in mice. Chromosoma 107, 570–576.

Kato, H., Jiang, J., Zhou, B.-R., Rozendaal, M., Feng, H., Ghirlando, R., Xiao, T.S., Straight, A.F., and Bai, Y. (2013). A conserved mechanism for centromeric nucleosome recognition by centromere protein CENP-C. Sci New York N Y 340, 1110–1113.

Katoh, K., and Standley, D.M. (2013). MAFFT Multiple Sequence Alignment Software Version 7: Improvements in Performance and Usability. Mol Biol Evol 30, 772–780.

Katoh, K., Misawa, K., Kuma, K., and Miyata, T. (2002). MAFFT: a novel method for rapid multiple sequence alignment based on fast Fourier transform. Nucleic Acids Res 30, 3059–3066.

Keane, T.M., Goodstadt, L., Danecek, P., White, M.A., Wong, K., Yalcin, B., Heger, A., Agam, A., Slater, G., Goodson, M., et al. (2011). Mouse genomic variation and its effect on phenotypes and gene regulation. Nature 477, 289–294.

Kelly, A.E., Ghenoiu, C., Xue, J.Z., Zierhut, C., Kimura, H., and Funabiki, H. (2010). Survivin Reads Phosphorylated Histone H3 Threonine 3 to Activate the Mitotic Kinase Aurora B. Science 330, 235–239.

Kim, J., Ishiguro, K., Nambu, A., Akiyoshi, B., Yokobayashi, S., Kagami, A., Ishiguro, T., Pendas, A.M., Takeda, N., Sakakibara, Y., et al. (2015). Meikin is a conserved regulator of meiosis-I-specific kinetochore function. Nature 517, 466–471.

Kipling, D., and Warburton, P.E. (1997). Centromeres, CENP-B and Tigger too. Trends Genet 13, 141–145.

Klare, K., Weir, J.R., Basilico, F., Zimniak, T., Massimiliano, L., Ludwigs, N., Herzog, F., and Musacchio, A. (2015). CENP-C is a blueprint for constitutive centromere-associated network assembly within human kinetochores. J Cell Biology 210, 11–22.

Krenn, V., Overlack, K., Primorac, I., van Gerwen, S., and Musacchio, A. (2013). KI motifs of human Knl1 enhance assembly of comprehensive spindle checkpoint complexes around MELT repeats. Curr Biology Cb 24, 29–39.

Lampson, M.A., and Black, B.E. (2017). Cellular and Molecular Mechanisms of Centromere Drive. Cold Spring Harb Sym 82, 249–257.

Langley, S.A., Miga, K.H., Karpen, G.H., and Langley, C.H. (2019). Haplotypes spanning centromeric regions reveal persistence of large blocks of archaic DNA. Elife 8, e42989.

Logsdon, G.A., Gambogi, C.W., Liskovykh, M.A., Barrey, E.J., Larionov, V., Miga, K.H., Heun, P., and Black, B.E. (2019). Human Artificial Chromosomes that Bypass Centromeric DNA. Cell 178, 624–639.e19.

Maheshwari, S., Tan, E.H., West, A., Franklin, F.C.H., Comai, L., and Chan, S.W.L. (2015). Naturally Occurring Differences in CENH3 Affect Chromosome Segregation in Zygotic Mitosis of Hybrids. Plos Genet 11, e1004970.

Malik, H.S., and Henikoff, S. (2001). Adaptive evolution of Cid, a centromere-specific histone in Drosophila. Genetics 157, 1293–1298.

Malik, H.S., and Henikoff, S. (2009). Major Evolutionary Transitions in Centromere Complexity. Cell 138, 1067–1082.

Masumoto, H., Masukata, H., Muro, Y., Nozaki, N., and Okazaki, T. (1989). A human centromere antigen (CENP-B) interacts with a short specific sequence in alphoid DNA, a human centromeric satellite. J Cell Biology 109, 1963–1973.

Melters, D.P., Bradnam, K.R., Young, H.A., Telis, N., May, M.R., Ruby, J.G., Sebra, R., Peluso, P., Eid, J., Rank, D., et al. (2013). Comparative analysis of tandem repeats from hundreds of species reveals unique insights into centromere evolution. Genome Biol 14, R10.

Moree, B., Meyer, C.B., Fuller, C.J., and Straight, A.F. (2011). CENP-C recruits M18BP1 to centromeres to promote CENP-A chromatin assembly. J Cell Biology 194, 855–871.

Musacchio, A., and Desai, A. (2017). A Molecular View of Kinetochore Assembly and Function. Biology 6, 5.

Nakagawa, H., Lee, J.-K., Hurwitz, J., Allshire, R.C., Nakayama, J., Grewal, S.I.S., Tanaka, K., and Murakami, Y. (2002). Fission yeast CENP-B homologs nucleate centromeric heterochromatin by promoting heterochromatin-specific histone tail modifications. Gene Dev 16, 1766–1778.

Nishino, T., Takeuchi, K., Gascoigne, K.E., Suzuki, A., Hori, T., Oyama, T., Morikawa, K., Cheeseman, I.M., and Fukagawa, T. (2012). CENP-T-W-S-X forms a unique centromeric chromatin structure with a histone-like fold. Cell 148, 487–501.

Nozawa, R.-S., Nagao, K., Masuda, H.-T., Iwasaki, O., Hirota, T., Nozaki, N., Kimura, H., and Obuse, C. (2010). Human POGZ modulates dissociation of HP1alpha from mitotic chromosome arms through Aurora B activation. Nat Cell Biol 12, 719–727.

Ohzeki, J.-I., Shono, N., Otake, K., Martins, N.M.C., Kugou, K., Kimura, H., Nagase, T., Larionov, V., Earnshaw, W.C., and Masumoto, H. (2016). KAT7/HBO1/MYST2 Regulates CENP-A Chromatin Assembly by Antagonizing Suv39h1-Mediated Centromere Inactivation. Dev Cell 37, 413–427.

Okada, T., Ohzeki, J., Nakano, M., Yoda, K., Brinkley, W.R., Larionov, V., and Masumoto, H. (2007). CENP-B Controls Centromere Formation Depending on the Chromatin Context. Cell 131, 1287–1300.

Otake, K., Ohzeki, J.-I., Shono, N., Kugou, K., Okazaki, K., Nagase, T., Yamakawa, H., Kouprina, N., Larionov, V., Kimura, H., et al. (2020). CENP-B creates alternative epigenetic chromatin states permissive for CENP-A or heterochromatin assembly. J Cell Sci jcs.243303.

Pacchierotti, F., Tiveron, C., Mailhes, J.B., and Davisson, M.T. (1995). Susceptibility to vinblastine-induced aneuploidy and preferential chromosome segregation during meiosis I in Robertsonian heterozygous mice. Teratogenesis Carcinog Mutagen 15, 217–230.

Pentakota, S., Zhou, K., Smith, C., Maffini, S., Petrovic, A., Morgan, G.P., Weir, J.R., Vetter, I.R., Musacchio, A., and Luger, K. (2017). Decoding the centromeric nucleosome through CENP-N. Elife 6, e33442.

Perelman, P., Johnson, W.E., Roos, C., Seuánez, H.N., Horvath, J.E., Moreira, M.A.M., Kessing, B., Pontius, J., Roelke, M., Rumpler, Y., et al. (2011). A molecular phylogeny of living primates. Plos Genet 7, e1001342.

Perez-Castro, A.V., Shamanski, F.L., Meneses, J.J., Lovato, T.L., Vogel, K.G., Moyzis, R.K., and Pedersen, R. (1998). Centromeric Protein B Null Mice Are Viable with No Apparent Abnormalities. Dev Biol 201, 135–143.

Pesenti, M.E., Prumbaum, D., Auckland, P., Smith, C.M., Faesen, A.C., Petrovic, A., Erent, M., Maffini, S., Pentakota, S., Weir, J.R., et al. (2018). Reconstitution of a 26-Subunit Human Kinetochore Reveals Cooperative Microtubule Binding by CENP-OPQUR and NDC80. Mol Cell 71, 923–939.e10.

Petrovic, A., Mosalaganti, S., Keller, J., Mattiuzzo, M., Overlack, K., Krenn, V., Antoni, A.D., Wohlgemuth, S., Cecatiello, V., Pasqualato, S., et al. (2014). Modular assembly of RWD domains on the Mis12 complex underlies outer kinetochore organization. Mol Cell 53, 591–605.

Petrovic, A., Keller, J., Liu, Y., Overlack, K., John, J., Dimitrova, Y.N., Jenni, S., Gerwen, S. van, Stege, P., Wohlgemuth, S., et al. (2016). Structure of the MIS12 Complex and Molecular Basis of Its Interaction with CENP-C at Human Kinetochores. Cell 167, 1028–1040.e15.

Rago, F., Gascoigne, K.E., and Cheeseman, I.M. (2015). Distinct Organization and Regulation of the Outer Kinetochore KMN Network Downstream of CENP-C and CENP-T. Curr Biol 25, 671–677.

Rizvi, S.M.A., Prajapati, H.K., and Ghosh, S.K. (2017). The 2 micron plasmid: a selfish genetic element with an optimized survival strategy within Saccharomyces cerevisiae. Curr Genet 64, 25–42.

Schueler, M.G., Swanson, W., Thomas, P.J., Program, N.C.S., and Green, E.D. (2010). Adaptive evolution of foundation kinetochore proteins in primates. Mol Biol Evol 27, 1585–1597.

Shi, L., Hu, E., Wang, Z., Liu, J., Li, J., Li, M., Chen, H., Yu, C., Jiang, T., and Su, B. (2016). Regional selection of the brain size regulating gene CASC5 provides new insight into human brain evolution. Hum Genet 136, 193–204.

Shono, N., Ohzeki, J., Otake, K., Martins, N.M.C., Nagase, T., Kimura, H., Larionov, V., Earnshaw, W.C., and Masumoto, H. (2015). CENP-C and CENP-I are key connecting factors for kinetochore and CENP-A assembly. J Cell Sci 128, 4572–4587.

Sironi, M., Cagliani, R., Forni, D., and Clerici, M. (2015). Evolutionary insights into host-pathogen interactions from mammalian sequence data. Nat Rev Genetics 16, 224–236.

Stamatakis, A. (2014). RAxML version 8: a tool for phylogenetic analysis and post-analysis of large phylogenies. Bioinformatics 30, 1312–1313.

Sugimoto, K., Kuriyama, K., Shibata, A., and Himeno, M. (1997). Characterization of internal DNA-binding and C-terminal dimerization domains of human centromere/kinetochore autoantigen CENP-C in vitro: role of DNA-binding and self-associating activities in kinetochore organization. Chromosome Res 5, 132–141.

Takeiri, A., Motoyama, S., Matsuzaki, K., Harada, A., Taketo, J., Katoh, C., Tanaka, K., and Mishima, M. (2013). New DNA probes to detect aneugenicity in rat bone marrow micronucleated cells by a pan-centromeric FISH analysis. Mutat Res 755, 73–80.

Talbert, P.B., Bryson, T.D., and Henikoff, S. (2004). Adaptive evolution of centromere proteins in plants and animals. J Biology 3, 18.

Thybert, D., Roller, M., Navarro, F.C.P., Fiddes, I., Streeter, I., Feig, C., Martin-Galvez, D., Kolmogorov, M., Janoušek, V., Akanni, W., et al. (2018). Repeat associated mechanisms of genome evolution and function revealed by theMus caroliandMus paharigenomes. Genome Res 28, 448–459.

Tromer, E., Snel, B., and Kops, G.J.P.L. (2015). Widespread Recurrent Patterns of Rapid Repeat Evolution in the Kinetochore Scaffold KNL1. Genome Biol Evol 7, 2383–2393.

Tsukahara, T., Tanno, Y., and Watanabe, Y. (2010). Phosphorylation of the CPC by Cdk1 promotes chromosome bi-orientation. Nature 467, 719–723.

van der Lee, R., Wiel, L., van Dam, T.J.P., and Huynen, M.A. (2017). Genome-scale detection of positive selection in nine primates predicts human-virus evolutionary conflicts. Nucleic Acids Res 45, 10634–10648.

Veld, P.J.H.I.’t, Jeganathan, S., Petrovic, A., Singh, P., John, J., Krenn, V., Weissmann, F., Bange, T., and Musacchio, A. (2016). Molecular basis of outer kinetochoreassembly on CENP-T. Elife 5, e21007.

Veld, P.J.H.I.’t, Volkov, V.A., Stender, I.D., Musacchio, A., and Dogterom, M. (2019). Molecular determinants of the Ska-Ndc80 interaction and their influence on microtubule tracking and force-coupling. Elife 8, e49539.

Wang, F., Dai, J., Daum, J.R., Niedzialkowska, E., Banerjee, B., Stukenberg, P.T., Gorbsky, G.J., and Higgins, J.M.G. (2010). Histone H3 Thr-3 Phosphorylation by Haspin Positions Aurora B at Centromeres in Mitosis. Science 330, 231–235.

Weir, J.R., Faesen, A.C., Klare, K., Petrovic, A., Basilico, F., Fischböck, J., Pentakota, S., Keller, J., Pesenti, M.E., Pan, D., et al. (2016). Insights from biochemical reconstitution into the architecture of human kinetochores. Nature 537, 249–253.

Yamagishi, Y., Honda, T., Tanno, Y., and Watanabe, Y. (2010). Two histone marks establish the inner centromere and chromosome bi-orientation. Sci New York N Y 330, 239–243.

Yan, K., Yang, J., Zhang, Z., McLaughlin, S.H., Chang, L., Fasci, D., Ehrenhofer-Murray, A.E., Heck, A.J.R., and Barford, D. (2019). Structure of the inner kinetochore CCAN complex assembled onto a centromeric nucleosome. Nature 574, 278–282.

Yang, Z. (2007). PAML 4: Phylogenetic Analysis by Maximum Likelihood. Mol Biol Evol 24, 1586–1591.

Yang, P., Wang, Y., and Macfarlan, T.S. (2017). The Role of KRAB-ZFPs in Transposable Element Repression and Mammalian Evolution. Trends Genetics Tig 33, 871–881.

Yao, Y., and Dai, W. (2012). Shugoshins function as a guardian for chromosomal stability in nuclear division. Cell Cycle Georget Tex 11, 2631–2642.

Zhao, G., Oztan, A., Ye, Y., and Schwarz, T.L. (2019). Kinetochore Proteins Have a Post-Mitotic Function in Neurodevelopment. Dev Cell 48, 873–882.e4.

